# *Kcns3* Deficiency Disrupts Parvalbumin Neuron Physiology in Mouse Prefrontal Cortex: Implications for the pathophysiology of schizophrenia

**DOI:** 10.1101/2020.09.02.280107

**Authors:** Takeaki Miyamae, Takanori Hashimoto, Monica Abraham, Rika Kawabata, Sho Koshikizawa, Yufan Bian, Mitsuru Kikuchi, G Bard Ermentrout, David A Lewis, Guillermo Gonzalez-Burgos

**Affiliations:** Translational Neuroscience Program, Department of Psychiatry, University of Pittsburgh School of Medicine, Pittsburgh PA, 15213, USA; Department of Psychiatry and Behavioral Science, Kanazawa University Graduate School of Medical Science, Kanazawa 920-8640, Japan; Research Center for Child Mental Development, Kanazawa University, Kanazawa 920-8640, Japan; Department of Mathematics, Faculty of Arts and Sciences, University of Pittsburgh, Pittsburgh PA, 15213

## Abstract

The unique fast spiking (FS) phenotype of cortical parvalbumin-positive (PV) neurons depends on multiple subtypes of voltage-gated potassium channels (Kv). PV neurons selectively express *Kcns3*, the gene encoding Kv9.3 subunits, suggesting that *Kcns3* expression is critical for the FS phenotype. *KCNS3* expression is lower in PV neurons in schizophrenia, but the effects of this alteration are unclear, because Kv9.3 subunit function is poorly understood. We therefore assessed the role of Kv9.3 subunits in PV neuron function by combining gene expression analyses, computational modeling, and electrophysiology in acute slices from the cortex of *Kcns3*-deficient mice

*Kcns3* mRNA levels were ~50% lower in cortical PV neurons from *Kcns3*-deficient relative to wildtype mice. While silent *per se*, Kv9.3 subunits are believed to amplify the Kv2.1 current in Kv2.1-Kv9.3 channel complexes. Hence, to assess the consequences of reducing Kv9.3 levels, we simulated the effects of decreasing the Kv2.1-mediated current in a computational model. The FS cell model with reduced Kv2.1 produced spike trains with irregular inter-spike intervals, or stuttering, and greater Na^+^ channel inactivation, possibly due to a smaller afterhyperpolarization. As in the computational model, PV basket cells (PVBCs) from *Kcns3*-deficient mice displayed spike trains with strong stuttering, which depressed PVBC firing, and smaller afterhyperpolarization. Moreover, *Kcns3* deficiency impaired the recruitment of PVBCs by stimuli mimicking synaptic input during cortical UP states, which elicited bursts of spikes at gamma frequency. Our data suggest that Kv9.3 subunits are critical for PVBC physiology, and that *KCNS3* deficiency in schizophrenia may impair the substrate of gamma oscillations.

**Significance statement:** In the neocortex, *Kcns3*, the gene encoding voltage-dependent potassium (Kv) channel subunits Kv9.3, is selectively expressed by parvalbumin-positive (PV) neurons. Moreover, *KCNS3* expression is decreased in PV neurons in schizophrenia. Kv 9.3 subunits are believed to amplify the current mediated by Kv2.1 subunits, however Kv9.3 function has not been investigated in PV cells.

Here, simulations in a computational model and electrophysiological experiments with *Kcns3*-deficient mice revealed that *Kcns3* deficiency disrupts repetitive firing in cortical PV neurons, possibly enhancing Na^+^ channel inactivation, and particularly with stimuli eliciting firing at gamma frequency band (30-80Hz). Our results suggest that Kv9.3 subunits are essential for PV neuron electrophysiology and that KCNS3 deficiency likely contributes to PV neuron dysfunction and gamma oscillation impairments in schizophrenia.

## Introduction

Parvalbumin-positive (PV) cells, a prominent subtype of cortical GABA neurons, shape pyramidal cell activity via inhibitory synaptic output that contributes to various network activity patterns, including gamma frequency oscillations (Tremblay et al., 2016). To produce this synaptic output, PV neurons can discharge trains of narrow spikes at high and constant frequency, displaying their classical fast spiking (FS) phenotype (Kawaguchi and Kubota, 1997; Hu et al., 2014).

The FS phenotype depends on the expression of voltage-gated K^+^ channel (Kv) subunits of the Kv3 subfamily (Du et al., 1996; Lau et al., 2000; Rudy and McBain, 2001). However, Kv3 expression is not sufficient to generate FS properties, since Kv3 subunits are also expressed by non-FS neurons (Bocksteins et al., 2012; Zhang et al., 2016; Paul et al., 2017), including pyramidal cells (Soares et al., 2017; Kelly et al., 2019). Moreover, FS properties likely depend on the combined effects of multiple Kv subtypes, as PV cells express multiple Kv subunit genes (Okaty et al., 2009; Paul et al., 2017; Enwright et al., 2018).

Previously, we assessed in the neocortex the expression of *KCNS3*, the gene encoding Kv9.3 subunits, and found that *KCNS3* transcription is PV neuron-selective (Georgiev et al., 2012), suggesting that Kv9.3 subunits play a crucial role in PV neuron electrophysiology. However, the function of Kv9.3 subunits is poorly understood (Bocksteins, 2016; Rasmussen and Trimmer, 2018) and has not been investigated in PV neurons.

Kv9.3 subunits are members of the silent subfamily of Kv subunits, which cannot form homomeric channels, and assemble into functional Kv channels only when combined with Kv2 subunits (Bocksteins, 2016; Rasmussen and Trimmer, 2018). Specifically, Kv9.3 subunits assemble with Kv2.1 to form Kv2.1-Kv9.3 heterotetramers (Bocksteins, 2016). PV neurons have the capacity to assemble Kv2.1-Kv9.3 channels, as they express *Kcnb1*, the gene encoding Kv2.1 subunits (Okaty et al., 2009; Paul et al., 2017; Enwright et al., 2018).

Kv2.1-Kv9.3 channels exhibit different biophysical properties than homomeric Kv2.1 channels, including ~two times greater single channel conductance (Patel et al., 1997) and more hyperpolarized voltage dependence of activation (Kerschensteiner and Stocker, 1999; Kerschensteiner et al., 2003). The biophysical properties of Kv2.1-Kv9.3 channels suggest that Kv9.3 subunits, while silent *per se*, amplify the Kv2.1-mediated K^+^ current (Bocksteins, 2016).

We reported that *KCNS3* mRNA levels are lower in PV neurons from the prefrontal cortex (PFC) of subjects with schizophrenia (Georgiev et al., 2014; Enwright et al., 2018). K^+^ channels produce hyperpolarizing currents that generally decrease neuronal excitability. Hence, one possibility is that *KCNS3* downregulation in schizophrenia contributes to a homeostatic increase in PV neuron excitability to restore PV neuron activity in a hypoactive network (Arion et al., 2015). However, Kv channels regulate excitability in complex ways (Rasmussen and Trimmer, 2018); thus, we suggested previously that *KCNS3* downregulation in schizophrenia could disrupt the time course of excitatory synaptic potentials in PV neurons (Georgiev et al., 2014). Understanding the role that Kv9.3 subunits play in PV neuron electrophysiology is therefore crucial to develop hypotheses regarding the effects of *KCNS3* deficiency on PV neurons and cortical circuit function in schizophrenia (Gonzalez-Burgos et al., 2015).

Using computational simulations and experiments with PV neurons from *Kcns3*-deficient mice, here we report that *Kcns3* deficiency disrupts the FS phenotype of PV basket cells (PVBCs), depressing recruitment of PVBCs at gamma band frequency by natural patterns of synaptic input. These effects of *Kcns3* deficiency suggest that Kv9.3 subunits play a crucial role in the transformation of synaptic input into spike output by PVBCs, and that lower *KCNS3* expression in schizophrenia could affect the recruitment of PV neurons by rhythmic excitatory input, possibly impairing the substrate of gamma oscillations.

## Materials and Methods

### Animals

Animal procedures followed the guidelines of the National Institutes of Health Guide for Care and Use of Laboratory Animals, and the Fundamental Guidelines for Proper Conduct of Animal Experiment and Related Activities in Academic Research Institutions from the Ministry of Education, Culture, Sports, Science and Technology of Japan, as approved by the IACUCs of the University of Pittsburgh and Kanazawa University.

#### Mouse lines

To create mice with deficiency in *Kcns3* expression, a trapping cassette consisting of LacZ-reporter and neomycin-resistance genes, each containing a polyadenylation signal sequence at the end, was inserted into the intron immediately upstream the only protein-coding exon of the *Kcns3* gene, by homologous recombination using the knockout first construct (CSD76140, Knockout Mouse Project Repository) to produce the *Kcns3*^tm1a(KOMP)Wtsi^ allele in C57BL/6-derived JM8 ES cells (Skarnes et al., 2011) (Figure 1A). The chimeric mice with a germline transmission of *Kcns3*^tm1a(KOMP)Wtsi^ allele were crossed with C57BL/6 mice to obtain heterozygous mice (MMRRC_065425-UCD).

**Figure 1.**
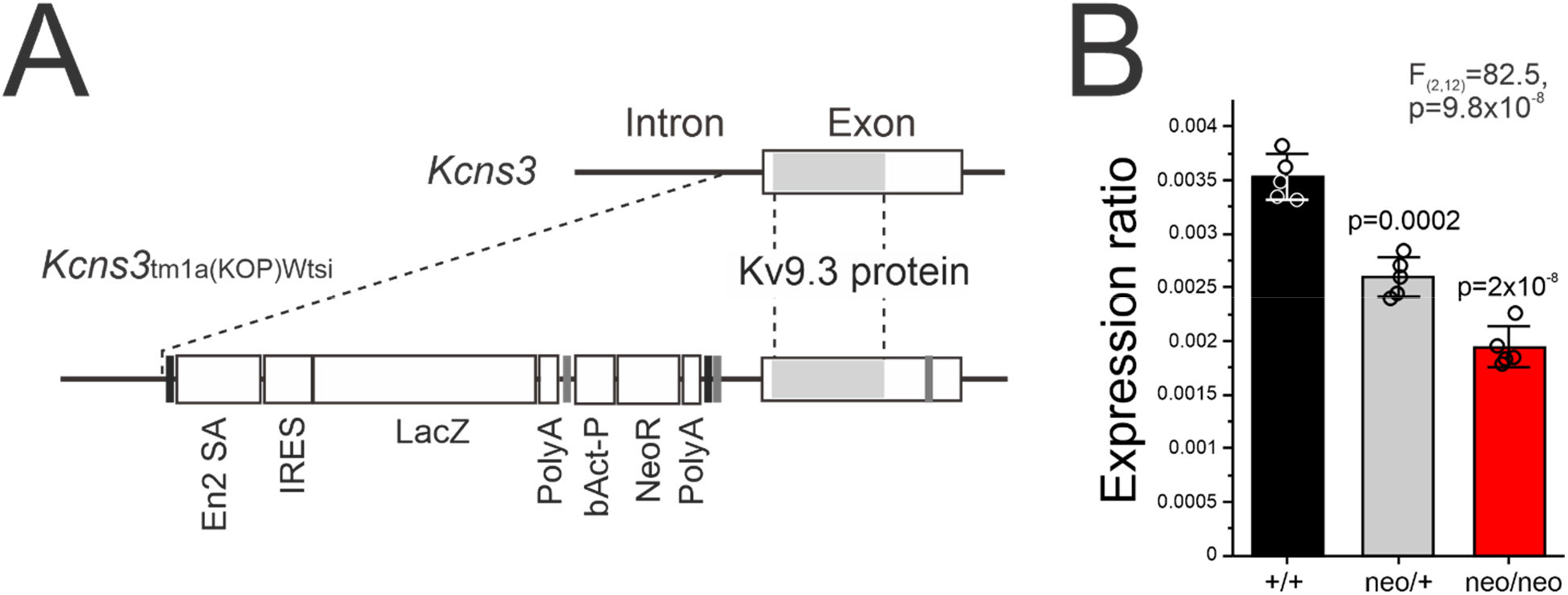
Gene trapping of *Kcns3.* **A)** The structure around the only protein-coding exon of wild type (top) and K*cns3*^tm1a(KOMP)Wtsi^ (bottom) alleles. The Kv9.3 protein-coding sequence in the exon is indicated by the shaded region. Broken lines connect the equivalent positions in both alleles. Vertical black lines: FRT sites, Vertical gray lines: loxP sites, En2SA: mouse En2 splicing acceptor site, IRES: Internal ribosomal entry site, LacZ; beta-galatosidase gene, Poly-A: SV40 polyadenylation sites, bAct-P: human *β*-actin promoter, NeoR: Neomycin resistance gene. **B)** Bar graphs summarizing the mean±SEM expression levels of *Kcns3* mRNA in the frontal cortex of *Kcns3*^+/+^, *Kcns3*^neo/+^ and *Kcns3*^neo/neo^ mice (n=5 per genotype) measured by real-time quantitative PCR (qPCR). Expression levels were ratios against the geometric mean of three internal control transcripts. The data of individual mice are presented by open circles superimposed over each bar. The F and p values reported are from One-way analysis of variance (ANOVA). P values of Tukey post-hoc comparisons versus *Kcns3^+/+^* mice are shown above each bar for *Kcns3*^neo/+^ and *Kcns3*^neo/neo^ mice.

G42 mice, which express green fluorescence protein (GFP) exclusively in PV neurons (Chattopadhyaya et al., 2004; Buchanan et al., 2012; Sippy and Yuste, 2013), were obtained from the Jackson Laboratory (stock No. 007677). The G42 line was generated by introducing a transgene consisting of regulatory and coding regions of the mouse *Gad1* gene in which the first protein-coding exon was replaced by cDNA for enhanced GFP (Chattopadhyaya et al., 2004).

#### Breeding

We crossed heterozygous *Kcns3*^tm1a(KOMP)Wtsi^ mice to obtain wild type (*Kcns3*^+/+^), heterozygous (*Kcns3*^neo/+^) and homozygous (*Kcns3*^neo/neo^) mice. In order to target PV neurons for electrophysiological recordings in acute slices, we crossed *Kcns3*^neo/+^ mice and G42^+/-^ mice (G42 mice that were heterozygous for the transgene). After crossing *Kcns3*^neo/+^ and G42^+/-^ mice, the resultant G42^+/-^;*Kcns3*^neo/+^ and G42^-/-^;*Kcns3*^neo/+^ mice were paired to produce *Kcns3^+/+^, Kcns3*^neo/+^ and *Kcns3*^neo/neo^ mice with G42^+/-^ or G42^-/-^ genotypes. Then, G42^+/-^; *Kcns3^+/+^* and G42^-/-^;*Kcns3*^+/+^ mice were bred to produce G42^+/-^;*Kcns3*^+/+^ mice, and G42^+/-^;*Kcns3*^neo/neo^ and G42^--^;*Kcns3*^neo/neo^ mice were bred to produce G42^+/-^;*Kcns3*^neo/neo^ mice. These G42^+/-^;*Kcns3^+/+^* and G42^+/-^;*Kcns3*^neo/neo^ mice are referred to as G42-*Kcns3*^+/+^ and G42-*Kcns3*^neo/neo^ mice, respectively, and were used as control and *Kcns3*-deficient animals in the electrophysiology experiments.

The mean±SD postnatal age of mice in six experimental groups of wild type and *Kcns3*-deficient mice was: 1) Gene expression analysis (Figures 1,2): 59±1.3 days for all genotypes; 2) Spontaneous excitatory postsynaptic potential (EPSP) recordings (Figure 4A-C): G42-*Kcns*3^+/+^: 32±2 days, G42-*Kcns3*^neo/neo^: 34±3 days; 3) Spontaneous excitatory postsynaptic current (EPSC) recordings (Figure 4D-F): G42-*Kcns*3^+/+^: 30±10 days, G42-*Kcns3*^neo/neo^: 27±4 days; 4) Intrinsic membrane properties (Figure 6): G42-*Kcns3*^+/+^: 38±11 days, G42-*Kcns3*^neo/neo^: 33±9 days; 5) Tests of the coefficient of variation of the inter spike intervals (CV_ISI_) (Figures 7 and 8): G42-*Kcns3*^+/+^: 40±8 days, G42-*Kcns3*^neo/neo^: 34±9 days; 6) Stimulation with UP states (Figures 9 and 10): G42-*Kcns3*^+/+^: 30±6 days, G42-*Kcns3*^neo/neo^: 32±6 days. Given that the postnatal development of PV neurons in the PFC of wild type mice is complete by postnatal day 21 (Miyamae et al., 2017), all PV neurons have adult-like electrophysiological properties at the age range used in the currents experiments.

**Figure 2.**
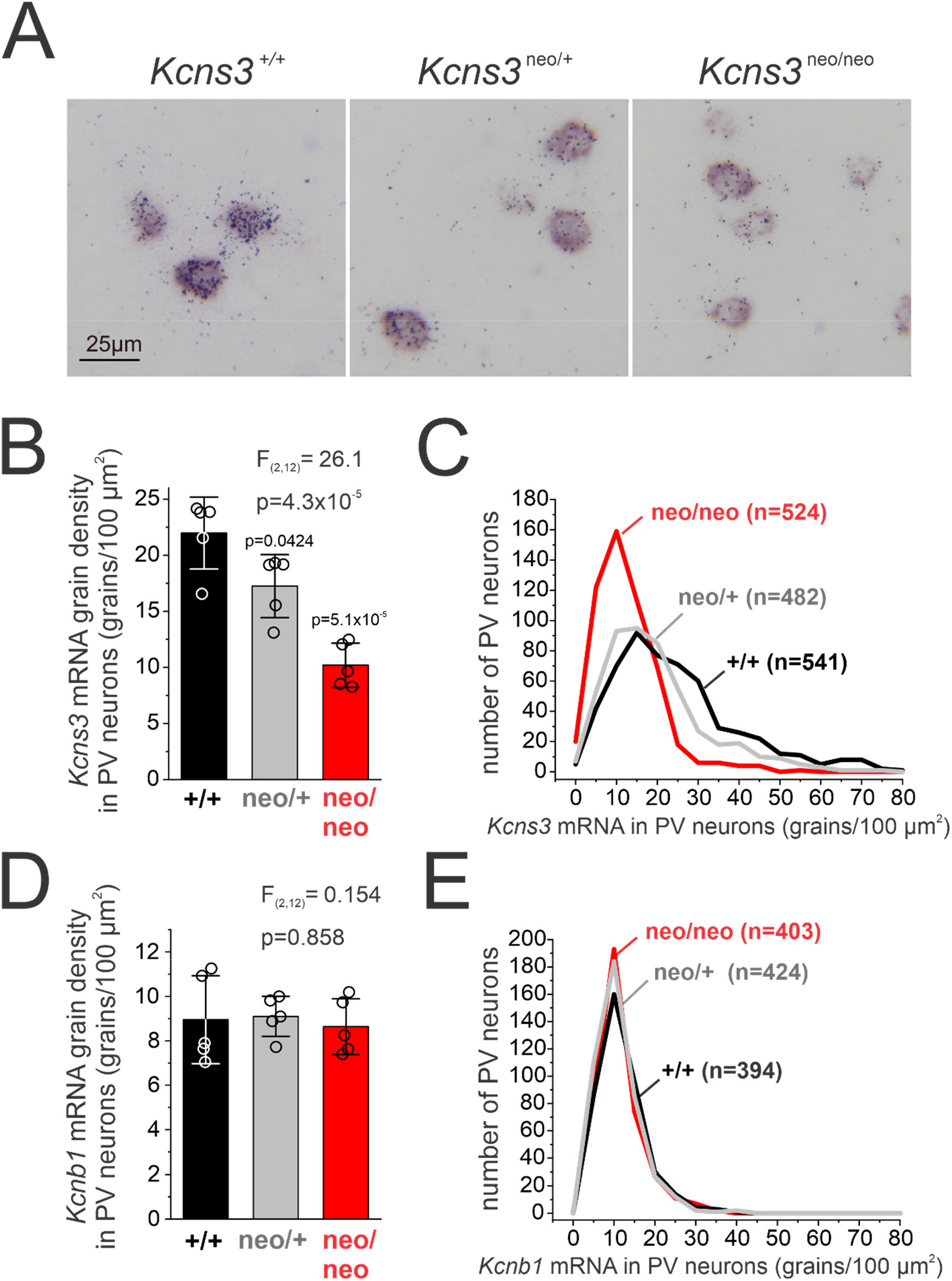
Gene expression in parvalbumin-positive (PV) neurons from *Kcns3*-deficient mice. **A)** Representative photomicrographs of frontal cortex from *Kcns3*^+/+^, *Kcns3*^neo/+^ and *Kcns3*^neo/neo^ mice showing expression of *Pvalb* mRNA (detected as the color reaction product by the digoxigenin-labeled riboprobe), and *Kcns3* mRNA (detected as silver grain accumulation by the ^35^S-labeled riboprobe). **B)** Bar graphs summarizing the mean±SEM *Kcns3* grain density per PV cell (grains/100 μm^2^) in frontal cortex of *Kcns3*^+/+^, *Kcns3*^neo/+^ and *Kcns3*^neo/neo^ mice (n=5, each genotype). The data of individual mice are presented by open circles superimposed over each bar. The F and p values reported are from One-way analysis of variance (ANOVA). P values of Tukey post-hoc comparisons versus*Kcns3^+/+^* mice are shown above each bar for *Kcns3*^neo/+^ and *Kcns3*^neo/neo^ mice. **C)** Histograms of distribution of *Kcns3* mRNA grain density per PV cell. **D)** Bar graphs summarizing the mean±SEM *Kcnb1* mRNA grain density per PV cell (grains/100μm^2^) in frontal cortex of *Kcns3*^+/+^, *Kcns3*^neo/+^ and *Kcns3*^neo/neo^ mice (n=5, each genotype). The data of individual mice are presented by open circles superimposed over each bar. The F and p values reported are from One-way ANOVA. **E)** Histograms of distribution of *Kcnb1* mRNA grain density per PV cell.

### Gene expression analysis

#### Quantitative PCR (qPCR)

Gray matter tissue was dissected from the frontal cortex at the level of +1.98mm to +1.70mm from bregma from *Kcns3*^+/+^, *Kcns3*^neo/+^, and *Kcns3*^neo/neo^ mice (n=5 per genotype) and homogenized in TRIzol reagent (Invitrogen, Carlsbad, CA). Total RNA samples were prepared from the homogenates using RNeasy lipid tissue mini kit (Qiagen, Hilden, Germany) and converted into cDNA with High Capacity RNA-to-cDNA Kit (Applied Biosystems, Foster City, CA). The *Kcns3* fragment was amplified with three internal control transcripts, betaactin *(Actb)*, cyclophilin A (Ppia) and glyceraldehyde-3-phosphate dehydrogenase (Gapdh), using Power SYBR Green PCR Master Mix and StepOnePlus real-time PCR system (Applied Biosystems). All primer sets (Table 1) amplified specific, single products with expected sizes and had amplification efficiency ≥96%. *Kcns3* mRNA levels were determined as 2^-dCT^ where dCT indicates the difference between cycle threshold (CT) of *Kcns3* and the mean CT of the three control transcripts.

**Table 1.**
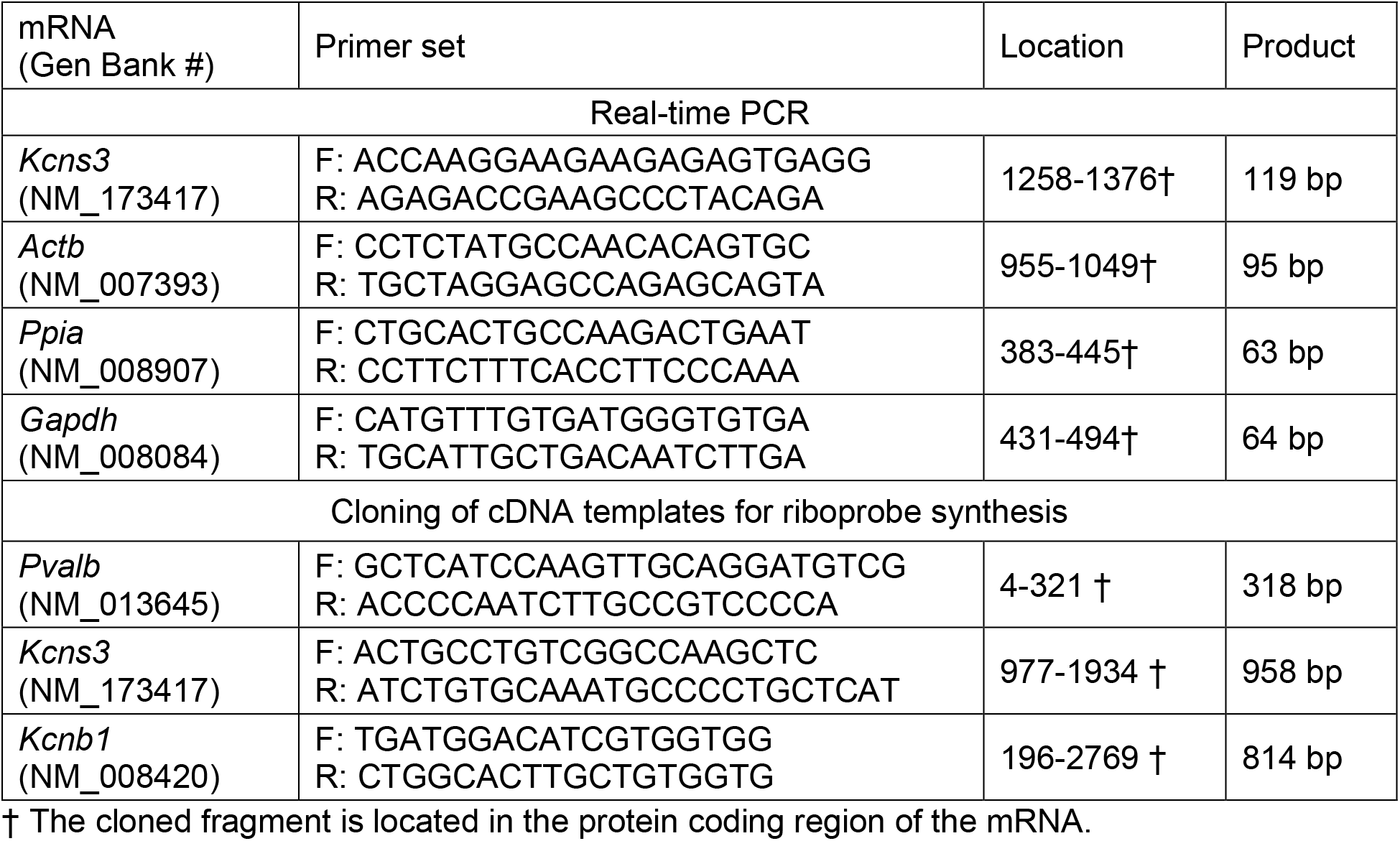
Primer sets used for real-time PCR and cloning of DNA templates.

#### In situ hybridization (ISH)

Coronal sections of the frontal cortex (12 μm) were cut from +1.98 mm to +1.54 mm from the bregma, and three sections evenly spaced at ≈130 μm intervals were selected from each mouse (n=5 per genotype) and subjected to ISH for each mRNA of interest. Templates cDNA fragments for the synthesis of riboprobes were obtained by PCR using specific primer sets (Table 1) and subcloned into the plasmid pSTBlue-1 (Novagen, Madison, WI). Nucleotide sequencing revealed 100% homologies of these fragments to the reported sequences in Genbank. Antisense and sense probes were transcribed *in vitro* in the presence of ^35^ S-CTP (PerkinElmer, Waltham, MA) for *Kcns3* and *Kcnb1* probes or in the presence of digoxigenin (DIG)-11-UTP (Roche, Mannheim, Germany) for *Pvalb* probe, using T7 or SP6 RNA polymerase (Promega, Madison, WI). All antisense riboprobes revealed distinctive signal distributions that were consistent with previous studies (Du et al., 1998; Georgiev et al., 2016) and Allen Brain Atlas (http://mouse.brain-map.org/). No signal beyond the background was detected with sense riboprobes for all mRNAs.

To detect *Kcns3* and *Kcnb1* mRNA levels in individual PV neurons, we performed dual-label ISH with ^35^S-labeled riboprobe for *Kcns3* or *Kcnb1* mRNA and digoxigenin (DIG)-labeled riboprobe for *Pvalb* mRNA. After fixation with 4% paraformaldehyde in phosphate-buffered saline (PBS), the sections were acetylated with 0.25% acetic anhydrate in 0.1 M triethanolamine/0.9% NaCl for 10 min and dehydrated through a graded ethanol series. The sections were then hybridized with ^35^-labeled riboprobes (2×10^7^ dpm/ml) for *Kcns3* or *Kcnb1* and DIG-labeled riboprobe (100 ng/ml) for *Pvalb* mRNA in hybridization buffer containing 50% formamide, 0.75 M NaCl, 20 mM 1,4-piperazine diethane sulfonic acid, pH 6.8, 10 mM EDTA, 10% dextran sulfate, 5× Denhardt’s solution (0.2 mg/ml Ficoll, 0.2 mg/ml polyvinylpyrrolidone, 0.2 mg/ml BSA), 50 mM dithiothreitol, 0.2% SDS, and 100 μg/ml yeast tRNA at 56°C for 16 hr. The sections were washed in a solution containing 0.3 M NaCl, 20 mM Tris-HCl, pH 8.0, 1mM EDTA, pH 8.0, and 50% formamide at 63°C, treated with 20 μg/ml RNase A (Sigma-Aldrich, St Louis, MO) at 37°C, and washed in 0.1× SSC (150 mM NaCl, 15 mM sodium citrate) at 66°C. After the washing, sections were preincubated in Tris-buffered saline (TBS, 100 mM Tris-HCl, pH 7.5, 150 mM NaCl) with 3% bovine serum albumin (BSA) and 0.05% Triton X-100 for 30 min, incubated with anti-DIG antibody conjugated with alkaline phosphatase (Roche) diluted 1:2000 in TBS with 1% BSA and 0.05% Triton X-100 for 12 hr at 4°C, washed in TBS, and air dried. To detect *Kcns3* or *Kcnb1* mRNA, sections were coated with NTB emulsion (Carestream, Rochester, NY) diluted 1:1 with distilled water at 43°C, exposed for 1 week at 4°C, and developed with D-19 (Carestream). For the detection of *Pvalb* mRNA, sections were incubated in 0.5 mg/ml nitroblue tetrazolium and 0.19 mg/ml bromo-chloroindolylphosphate (Roche) in 100 mM Tris-HCl, pH 9.5, 100 mM NaCl, 50 mM MgCl2 for 24 hr.

#### Quantification of gene expression in individual PV neurons

For quantification of *Kcns3* and *Kcnb1* mRNA levels in PV neurons, the density of silver grains representing the expression of these mRNAs was determined in individual PV neurons in the frontal cortex of *Kcns3*^+/+^, *Kcns3*^neo/+^ and *Kcns3*^neo/neo^ mice (n=5 per genotype). Using a microscope equipped with a motor-driven stage (Prior Scientific, Cambridge, UK) and the microcomputer imaging device (MCID) system (InterFocus Imaging, Cambridge, UK), sampling frames with a size of 200 × 140 μm were placed systematically and randomly in the region of interest (ROI) over the frontal cortex that included motor and primary sensory regions. In each frame, we counted the number of silver grains within the contour of individual PV neurons defined by alkaline phosphatase color reaction for *Pvalb* mRNA expression and calculated the raw density of silver grains per 100 μm^2^ PV neuron area. Three sections were analyzed for each mouse. In each section, mean background grain density (per 100 μm^2^) was determined in gray matter regions devoid of a grain cluster or color reaction. The density of silver grains in individual PV neurons was determined by subtracting the background grain density from the raw grain density of individual PV neurons sampled in the same section. For *Kcns3* mRNA, 108±15, 96±8.3 and 104±17 PV neurons per mouse were analyzed in *Kcns3*^+/+^, *Kcns3*^neo/+^ and *Kcns3*^neo/neo^ mice, respectively. For *Kcnb1* mRNA, 79±18, 85±18 and 81±16 PV neurons per mouse were analyzed in *Kcns3^+/+^, Kcns3*^neo/+^ and *Kcns3^neo/neo^* mice, respectively.

### Brain slice preparation and electrophysiology

#### Solutions employed

Slicing solution (in mM): sucrose 210, NaCl 10, KCl 1.9, Na_2_HPO_4_ 1.2, NaHCO_3_, MgCl_2_ 6, CaCl_2_ 0.5, glucose 10, pH 7.3–7.4 when bubbled with 95% O_2_-5% CO_2_.

Artificial Cerebro Spinal Fluid (ACSF, in mM): NaCl 125, KCl 2.5, Na_2_HPO_4_ 1.25, glucose 10, NaHCO3 25, MgCl_2_ 1 and CaCl_2_ 2, pH 7.3–7.4, bubbled with 95% O_2_-5% CO_2_.

ACSF modified to induce UP states (in mM): NaCl 125, KCl 3.5, Na_2_HPO_4_ 1.25, glucose 10, NaHCO_3_ 25, MgCl_2_ 1 and CaCl_2_ 1, pH 7.3–7.4, bubbled with 95% O_2_-5% CO_2_.

Potassium Gluconate (in mM): KGluconate 120, KCl 10, HEPES 10, EGTA 0.2, MgATP 4.5, NaGTP 0.3, sodium phosphocreatine 14, pH 7.2–7.4.

### Slice preparation

Brain slice preparation was performed as described previously (Miyamae et al., 2017). Briefly, coronal slices (300 μm) were cut from the frontal cortex in ice-cold slicing solution. After cutting, slices were incubated at room temperature in ACSF. For recording, slices were submerged in a chamber superfused with oxygenated ACSF (2-3 ml/min, 30-32°C) in the presence of the GABA_A_ receptor antagonist gabazine (10 μM) when recording sEPSPs or sEPSCs. To record intrinsic membrane properties and during UP state stimulation, the AMPA receptor antagonist cyanquixaline (CNQX) (10 μM) was added. Current clamp recordings were obtained from PV neurons identified by GFP fluorescence in layers 2 to 5, using microscopes (Zeiss Axioskop FS; Olympus BX51) equipped with epifluorescence, infrared illumination, differential interference contrast, and CCD video cameras (EXi Aqua, Q-Imaging, AZ). Glass pipettes (3-5 MΩ) were filled with Potassium Gluconate solution containing biocytin (0.4%) freshly added to the pipette solution for later identification of the PV cell morphology. For whole-cell recordings, we used Multiclamp 700A/B amplifiers (Axon Instruments, CA) with low-pass filtering at 4 kHz, and digitizing at 10/20 kHz using Power 1401 digital-analog converter and the Signal 5 software (Cambridge Electronic Design, CED, Cambridge, UK). After recordings, the slices were immersed in 4% paraformaldehyde in 0.1 M PBS for 24-72 h at 4°C and were subject to diaminobenzidine-based processing to visualize the biocytin for reconstruction of the morphology of the recorded neurons using Neurolucida (MBF Bioscience, VT) (Miyamae et al., 2017).

### Spontaneous excitatory postsynaptic potentials (sEPSPs) and excitatory postsynaptic currents (sEPSCs)

#### sEPSPs

Current clamp recordings were performed as described previously (Miyamae et al., 2017). The series resistance and pipette capacitance were monitored and cancelled using bridge balance and capacitance neutralization circuits in the Multiclamp amplifiers (Axon Instruments). We recorded sEPSPs in the presence of gabazine (10 μM) at the cell’s resting membrane potential (RMP) for 3-5 min, and 100-200 sEPSPs were detected for each PV neuron using MiniAnalysis software (Synaptosoft, GA). For the analysis of large sEPSPs from the recording, we selected all the sEPSPs with peak amplitude larger than a threshold of 1 mV. The sEPSP amplitude reported for each neuron is the average of the amplitudes of all the sEPSPs detected in each cell. The decay kinetics was quantified on the average sEPSP waveform obtained for each cell, and fitting a double exponential decay function and computing a weighted decay time constant (τw) as follows (Rotaru et al., 2011):

τw=[(A_slow_ x τ_slow_)+(A_fast_ x τ_fast_)]/(A_slow_ + A_fast_), where A_slow_, A_fast_, τ_slow_, τ_fast_ are the amplitude and time constants of slow and fast EPSP decay components.

Tetraethylammonium (TEA) was dissolved in ACSF at the final concentration (10 mM) used in the experiments and bath-applied for 10 minutes. TEA and all other chemical reagents were obtained from Sigma-Aldrich (MO).

#### sEPSCs

Voltage clamp recordings were performed as described previously (Miyamae et al., 2017). sEPSCs were recorded at the holding potential of −70mV, near the RMP. The stability of the series resistance (Rs) was monitored by the current evoked by a 50ms, +10mV step, delivered every 10s. Only recordings with an initial Rs <15 MΩ were used for analysis as described in detail previously (Miyamae et al., 2017). The measurements of sEPSC amplitude and decay kinetics were as described for sEPSPs above.

#### Injection of rectangular current steps

During current injection experiments, the series resistance and pipette capacitance were monitored and cancelled using bridge balance and capacitance neutralization circuits in the Multiclamp amplifiers (Axon Instruments). Families of rectangular 500 ms current pulses were applied in 10 pA increments, starting at −50 pA and repeating each step amplitude 2-3 times at 0.2 Hz. The following intrinsic membrane properties were measured from each PV cell: RMP, action potential (AP) voltage threshold, AP current threshold or rheobase, AP amplitude, AP half width, first AP delay, afterhyperpolarization, input resistance (Rin), slope of the frequency-current plot (f-I slope) and membrane time constant, as described previously (Miyamae et al., 2017). The rheobase was estimated as the smallest depolarizing current step amplitude that elicited at least one AP in each repetition of that particular step amplitude. Spike frequency adaptation was estimated calculating the ratio between the last and first inter-spike intervals (ISIs) in each spike train, or ISI_ratio_ = ISI(n)/ISI(1). The variability of firing was measured by the coefficient of variation of the ISIs (CV_ISI_), i.e. the standard deviation (SD) of the ISIs divided by their mean. The CV_ISI_ was estimated in the spike trains evoked by each current step amplitude. The analysis of ISIs was performed using scripts written in Signal 5 (CED) and OriginPro (OriginLab, MA) software packages. For the CV_ISI_, the data are expressed as a function of current above the threshold level that produced at least three spikes, which in many individual cells was higher than the rheobase.

#### Injection of current mimicking UP states

To assess the effects of *Kcns3* deficiency on the spike output elicited by natural patterns of stimulus, we injected into PV neurons depolarizing current mimicking the time course and amplitude of synaptic input observed during UP states. First, UP states were induced in acute slices from the PFC of G42 mice by perfusing the recording chamber with modified ACSF at a high flow rate (8-12 ml/min) (Sanchez-Vives and McCormick, 2000). Under these conditions, UP states developed in PFC cells within a few minutes of the perfusion (Sanchez-Vives and McCormick, 2000). Then, synaptic input during a sequence of 3 subthreshold UP states (UP_(1)_, UP_(2)_ and UP_(3)_) was recorded from an individual neuron and digitally stored. The time course and amplitude pattern of synaptic input during these 3 UP states was used in current clamp experiments to inject depolarizing current into PV neurons via the recording electrode, reproducing the changes in membrane potential observed during the UP states.

As the recorded UP states were subthreshold, firing was elicited by increasing the mean current injected during UP_(1)_ in increments of 10 pA, which resulted in increments of 7 and 8 pA of the mean current injected during UP_(2)_ and UP_(3)_, respectively. The smallest injected current that elicited spikes during UP_(1)_was defined as the stimulus strength 1, or threshold. Thereafter, 3 additional increments were applied to a total of 4 stimulus strengths. Each stimulus strength was repeated in 10 trials (intertrial interval=5 s). The spike probability, p(spike), was estimated across the duration of each UP state in 2 ms bins. The p(spike) was calculated as: p(spike)= number of trials with spikes/10, where 10 is the total number of trials per stimulus strength. The analysis of spike output during UP state stimulation was performed using scripts written in Signal 5 (CED) and OriginPro (OriginLab) software packages.

### Computational modeling

A Hodgkin-Huxley type computer model of a fast-spiking (FS) neuron was implemented using XPP software (http://www.math.pitt.edu/~bard/xpp/xpp.html). The model we used for the simulations was based on Erisir et al., 1999 (Erisir et al., 1999) with several modifications. First, we included a slow inactivation of the sodium current based on equations published by Fleidervish and colleagues (Fleidervish et al., 1996). We added an additional potassium conductance to model the Kv2.1 channel, g_Kv2_. We set the slow potassium channel conductance to 0 to eliminate subthreshold oscillations. We added a Gaussian white noise term to the voltage to mimic intrinsic noise in the physiological preparation. For completeness, below we include the full set of equations:

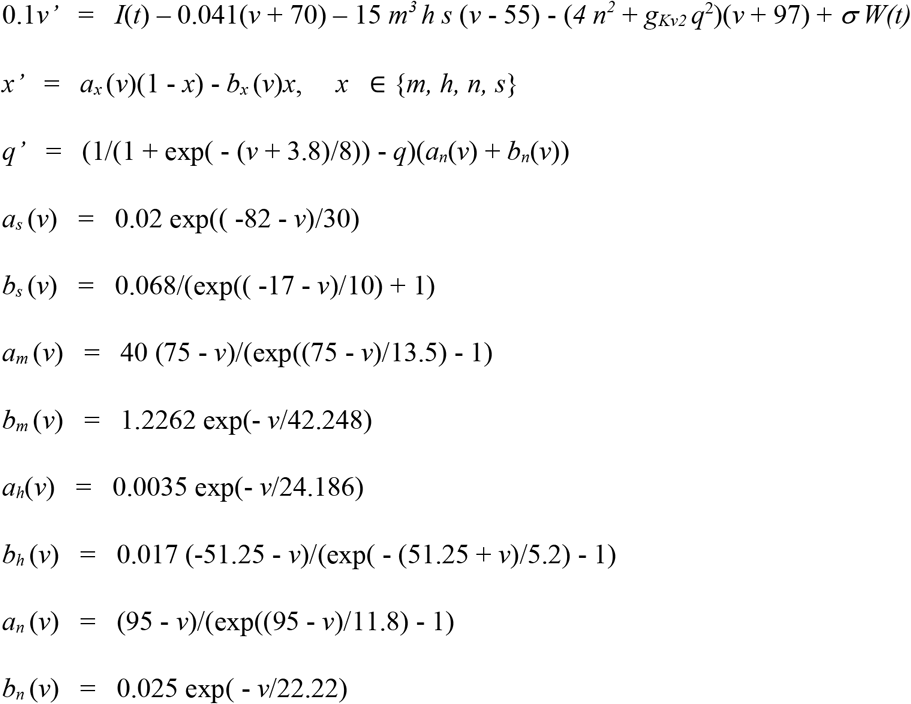

I(t) is a step function of current with various values as given in Figure 5. Similarly, the noise amplitude, σ, and the Kv2 conductance, g_Kv2_, also varied. W(t) is a white noise increment. *m, h*and *s* are the activation (m) and inactivation (h, s) gating variables for the Na^+^ conductance, and *n* and *q* are the activation gate variables for the Kv conductances.

### Statistical analysis

The data are expressed as mean±SD unless otherwise indicated. Analysis was performed using SPSS 25 (IBM, NY). ISH data: One-way ANOVA followed by Tukey’s test were used to compare *Kcns3* and *Kcnb1* mRNA levels in PV neurons across genotypes. Electrophysiology data: The normality of data distribution was tested on the residuals of the data using Shapiro Wilk tests and was verified by using the D’Agostino K-squared test, in every case both normality tests produced identical results. When Shapiro-Wilk tests implemented on the residuals of the data rejected normality, the tests were repeated on the residuals of the Ln-transformed data (Miyamae et al., 2017). The differences between groups were assessed using Student’s t-test and paired samples t-test. Nonparametric Mann-Whitney’s or Wilcoxon Signed Rank tests were performed on the original data if normality was rejected after Ln transformation. Shapiro Wilk tests with p<0.05 are indicated in the Figure legends.

## Results

### In *Kcns3^neo/neo^* mice, PV neurons show *Kcns3* downregulation with unaltered *Kcnb1*expression

To assess the extent to which *Kcns3* expression is affected by the insertion of the trapping cassette (Figure 1A), we performed qPCR using RNA isolated from the frontal cortex (Figure 1B) and found that relative to *Kcns3*^+/+^ mice, *Kcns3* mRNA levels were lower by 26% and 45% in *Kcns3*^neo/+^ and *Kcns3*^neo/neo^ mice, respectively (One-Way ANOVA, F(2,12)=82.5, p=9.8×10^-8^).

Next, we examined the effects of the genetic manipulation on *Kcns3* expression in individual PV neurons. We performed dual label-ISH in the frontal cortex from *Kcns3*^+/+^, *Kcns3*^neo/+^ and *Kcns3*^neo/neo^ mice (Figure 2A), quantifying *Kcns3* mRNA grain density in *Pvalb*mRNA-positive cells. The mean density of *Kcns3* mRNA grains per PV neuron was decreased by 21% and 53% in *Kcns3*^neo/+^ and *Kcns3*^neo/neo^ mice, respectively, relative to *Kcns3^+/+^* mice (Figure 2B). To assess the heterogeneity of the decrease in *Kcns3* gene expression per cell, we examined the distribution of *Kcns3* mRNA grain densities per individual PV neuron, and found that the distributions were left-shifted in both *Kcns3*^neo/neo^ and *Kcns3*^neo/+^ relative to *Kcns3*^+/+^ mice (Figure 2C).

Given that the main role hypothesized for Kv9.3 subunits is amplifying the Kv2.1 current, one possibility is that *Kcns3* deficiency, by decreasing Kv2.1 amplification, causes compensatory upregulation of the expression of *Kcnb1*, the gene encoding Kv2.1 subunits that is ubiquitously expressed by a large fraction cortical PV neurons (Du et al., 1998) (Allen Brain Atlas, https://celltypes.brain-map.org/rnaseq#transcriptomics). However, we found that *Kcnb1*mRNA levels in PV neurons did not differ among *Kcns3^+/+^*, *Kcns3*^neo/+^ and *Kcns3*^neo/neo^ mice (Figure 2D, E), suggesting an absence of compensatory upregulation of Kv2.1 subunits in response to *Kcns3* deficiency. Thus, the main effect of *Kcns3* deficiency is likely to be a decrease of Kv2.1 current caused by a partial loss of Kv9.3-mediated amplification.

### *Kcns3* deficiency does not affect excitatory synaptic inputs onto PV basket cells

PV neuron physiology is tuned for fast input-output transformation (Jonas et al., 2004; Hu et al., 2014), displaying rapid synaptic events at their excitatory inputs (Geiger et al., 1997; Angulo et al., 1999; Rotaru et al., 2011). Previously, we hypothesized that, in PV neurons, Kv2.1-Kv9.3 channel activation by subthreshold EPSPs elicits a K^+^ current that accelerates the EPSP decay, and hence *Kcns3* deficiency prolongs the EPSP decay (Georgiev et al., 2012; Georgiev et al., 2014).

To test the hypothesis that *Kcns3* deficiency prolongs the EPSP decay, we obtained current clamp recordings from layers 2-5 PV neurons in slices from the PFC of G42-*Kcns3*^+/+^ and G42-*Kcns3*^neo/neo^ mice. The recorded PV neurons were filled with biocytin to assess their morphology after the experiments. Neuronal reconstructions showed that the great majority of recorded PV neurons (92.5%) were basket cells (PVBCs), as illustrated by the examples in Figure 3A,B. Moreover, all cells contributing data to Figures 4, 6-10 (G42-*Kcns3^+/+^*, n=93; G42-*Kcns3*^neo/neo^, n=80) were PVBCs, and only a minority (7.5%) of the recorded PV neurons (G42-*Kcns3*^+/+^, n=5 cells; G42-*Kcns3*^neo/neo^, n=8 cells) were chandelier cells (data not shown), which, as in our previous studies (Miyamae et al., 2017; Tikhonova et al., 2018), had cell bodies exclusively localized near the border between layers 1 and 2.

**Figure 3.**
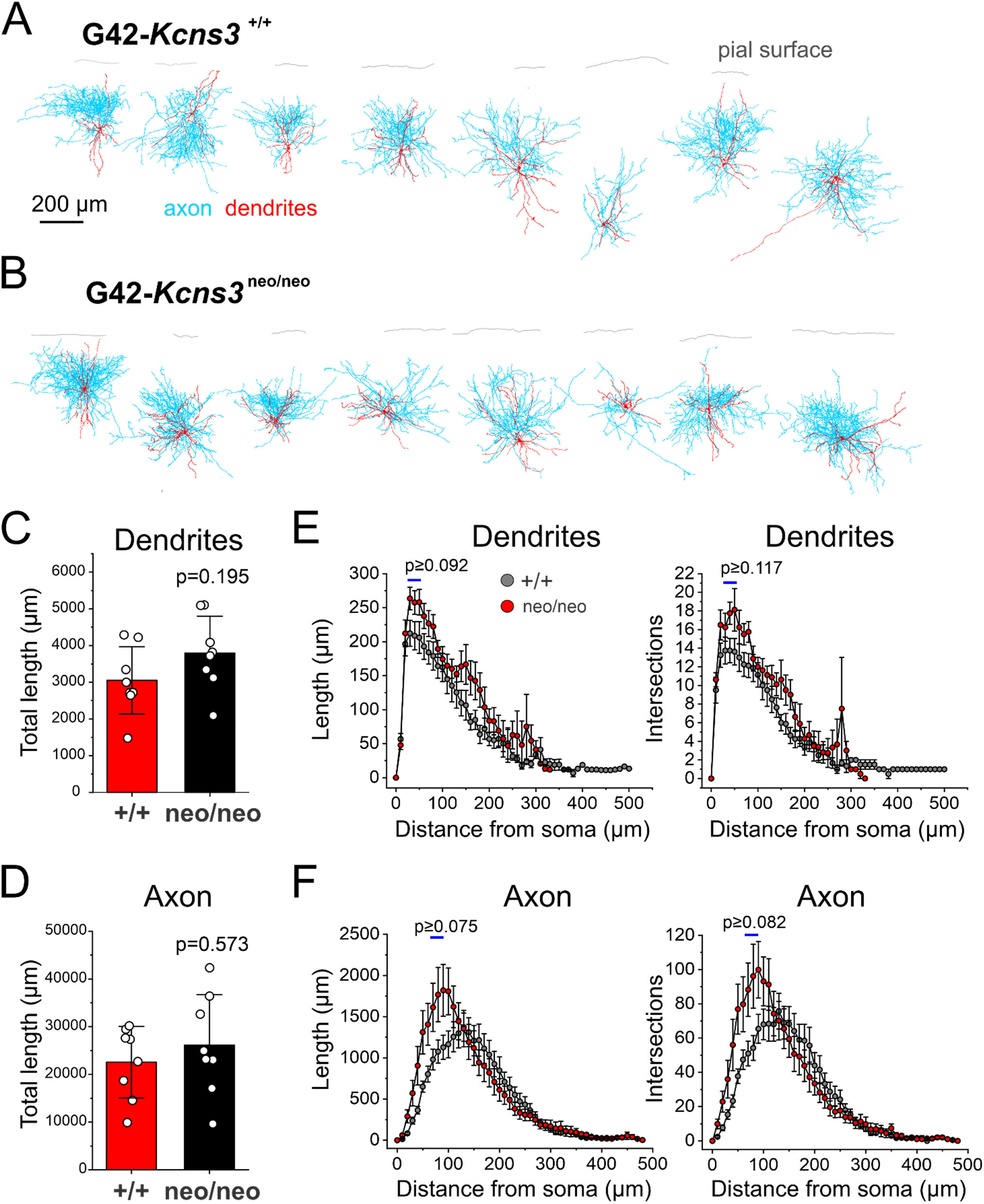
Morphology of the parvalbumin-positive (PV) neurons studied in prefrontal cortex slices from G42-*Kcns3*^+/+^ and G42-*Kcns3*^neo/neo^ mice. **A)** Digital reconstructions (n=8) of PV basket cells (PVBCs) recorded in slices from G42-*Kcns3*^+/+^ mice. **B)** Digital reconstructions (n=8) of PVBCs recorded in slices from G42-*Kcns3*^neo/neo^ mice. Calibration bar and labels apply to both A) and B). **C)** Bar graph displaying the total length of the dendritic tree of PVBCs from G42-*Kcns3*^+/+^ (n=8 cells from 5 mice) and G42-*Kcns3*^neo/neo^ mice (n=8 cells from 5 mice). Shown here, and in part D), are the p values from Mann-Whitney tests performed after Shapiro-Wilk tests of the residuals produced p<0.05 for both the original data and the Ln-transformed data. **D)** Bar graph displaying the total length of the axonal arbor of PVBCs from G42-*Kcns3*^+/+^ and G42-*Kcns3*^neo/neo^ mice, same cells as in C). **E)** Plots of mean±SEM for dendrite length (left panel) and number of intersections (right panel) from Sholl analysis of the same neurons in C). Shown here, and in panel F), are the smallest p values from t-test comparisons done for the three Sholl compartments with largest differences between genotypes, indicated by horizontal bars. **F)** Plots of mean±SEM for axon length (left panel) and number of intersections (right panel) from Sholl analysis of the same neurons in C). The p values are from t-test comparisons, as indicated in E).

In the PVBCs with digital reconstructions (Figure 3A,B), quantitative morphometry analysis (G42-*Kcns3^+/+^* n=8 cells from 5 mice; G42-*Kcns3*^neo/neo^, n=8 cells from 5 mice) showed that neither the total length of the dendritic tree (Figure 3C) nor axonal arbor (Figure 3D) differed by genotype. Sholl analysis showed a trend for greater length and branching complexity in the proximal compartments of the dendrites and axon of PVBCs from *Kcns3*-deficient mice (Figure 3E,F), but these differences did not achieve statistical significance.

PVBCs showed frequent sEPSPs with a wide range of peak amplitudes and fast decay (Figure 4A). However, contrary to our predictions, the decay time course of the average sEPSP did not differ by genotype (Figure 4B). The absence of effect of *Kcns3* deficiency on the sEPSP decay phase was not due to absence of regulation of the sEPSP decay by Kv channels, because bath application of 10 mM tetraethylammonium (TEA), a non-selective Kv channel blocker, prolonged the sEPSP decay time in PVBCs from G42-*Kcns3*^+/+^ mice (control: 6.8±2.4 ms, TEA: 8.5±3.2 ms, n=8 cells from 2 mice, p=0.0265, paired samples t-test), as reported for hippocampal PVBCs (Hu et al., 2010).

**Figure 4.**
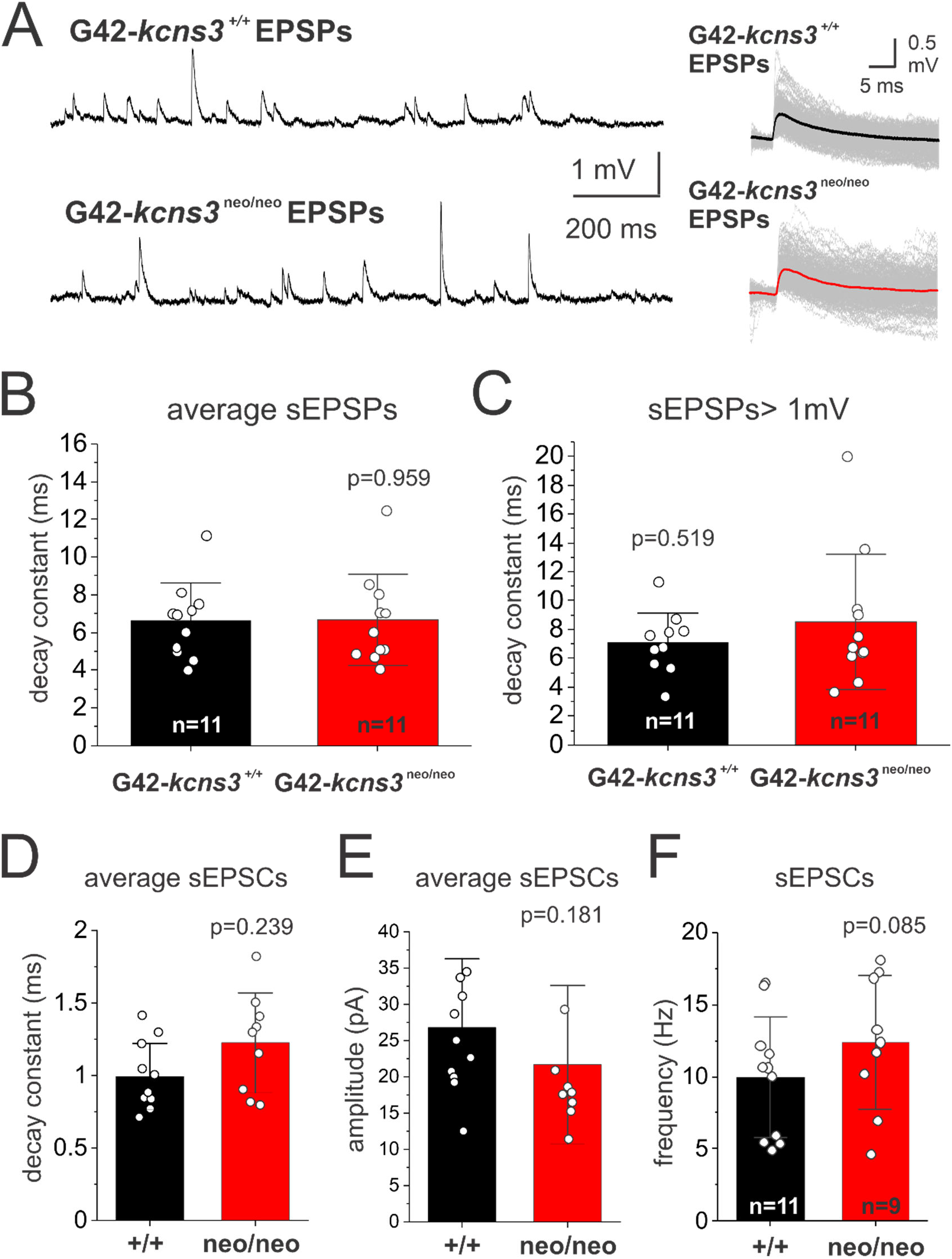
Effects of *Kcns3* deficiency on excitatory inputs to parvalbumin-positive basket cells (PVBCs). **A)** Left panel, traces illustrating examples of sEPSPs recorded at resting membrane potential (RMP) from PVBCs in slices from G42-*Kcns3*^+/+^ and G42-*Kcns3*^neo/neo^ mice. Right panel, examples of individual sEPSPs recorded from PVBCs of G42-*Kcns3*^+/+^ and G42-*Kcns3*^neo/neo^ mice, aligned by their rising phase and superimposed with their averages (thick lines). **B)** Bar graphs illustrating the decay time constant of sEPSPs recorded from PVBCs from G42-*Kcns3*^+/+^ and G42-*Kcns3*^neo/neo^ mice. **C)** Bar graphs illustrating the decay time constant of sEPSPs> 1 mV recorded from PVBCs from G42-*Kcns3*^+/+^ and G42-*Kcns3*^neo/neo^ mice. **D) E)** and **F)** Bar graphs illustrating the decay time constant, peak amplitude and frequency of sEPSCs recorded from PVBCs from G42-*Kcns3*^+/+^ and G42-*Kcns3*^neo/neo^ mice. In B)-E), shown are the p values of comparisons performed using Mann Whitney’s test, as the Shapiro-Wilk analysis of the residuals produced p<0.05 for both the original data and the Ln-transformed data.

It is possible that only the sEPSPs with largest amplitudes, which are a small fraction of all sEPSPs (Figure 4A), produce enough depolarization to activate the Kv2.1-Kv9.3 channels and accelerate the sEPSP decay time course. If so, the large sEPSPs should display longer decay phase in PVBCs from *Kcns3*^neo/neo^ versus *Kcns3^+/+^* mice. To test this prediction, we selected the sEPSPs with peak amplitude >1 mV (sEPSPs>1 mV). The sEPSPs>1 mV had peak amplitude ~3 times greater than the average sEPSP (G42-*kcns3^+/+^:*1.5 ± 0.3 mV versus 0.6 ± 0.2 mV, n=11, p= 0.000081, Mann Whitney’s test; G42-*kcns3*^neo/neo^: 1.9±1.1 mV versus 0.6 ± 0.1 mV, n=11; p= 0.0000028, Mann Whitney’s test). However, the decay time constant of the sEPSPs>1 mV did not differ between genotypes (Figure 4C).

*Kcns3* deficiency may change the properties of excitatory postsynaptic currents (EPSCs) in PVBCs indirectly, for example, secondary to changes in network activity that modify synaptic strength. To test whether *Kcns3* deficiency affects the excitatory input onto PVBCs, we assessed the properties of sEPSCs recorded from PVBCs in the PFC of *Kcns3*^neo/neo^ (n=9 cells from 4 mice) or *Kcns3^+/+^* mice (n=11 cells from 4 mice). Neither the sEPSC decay time constant (Figure 4D), nor the sEPSC amplitude (Figure 4E) were significantly different in PVBCs from *Kcns3*-deficient mice relative to *Kcns3^+/+^* mice. Moreover, the sEPSC frequency did not differ between *Kcns3*^neo/neo^ and *Kcns3*^+/+^ mice (Figure 4F), thus additionally suggesting absence of change in the excitatory drive onto PVBCs from *Kcns3-*deficient mice.

### Kv2.1 deficiency alters repetitive firing in a computational model of FS neurons

Previous studies suggested that Kv2.1 channels regulate repetitive firing. For example, in layer 3 pyramidal cells and spinal motoneurons, Kv2.1 currents facilitate repetitive firing by preventing Na^+^ channel inactivation (Guan et al., 2013; Romer et al., 2019). The FS phenotype of PV neurons, however, differs significantly from the firing patterns of pyramidal cells or motoneurons. Hence, to prove the concept that Kv2.1 currents regulate repetitive firing in PV neurons, we used a computational model of FS cells to simulate the effects of reducing the Kv2.1 current, which is the expected consequence from a loss of Kv9.3-mediated amplification of Kv2.1 current with *Kcns3* deficiency. To assess the effects of reducing Kv2.1 current, we used a well-known Hodgkin-Huxley type model of FS cells (Erisir et al., 1999), modified by adding the Kv conductance g_Kv2_, to simulate the Kv2.1 current, and a slow inactivation component to the voltage-gated Na^+^ conductance (Martina and Jonas, 1997) using previously published parameters (Fleidervish et al., 1996). With g_Kv2_ intact, application of excitatory current to the FS cell model produced spike trains with the continuous firing typical of the FS phenotype (Goldberg et al., 2008) (Figure 5A). In contrast, in the FS cell model with reduced g_Kv2_ levels, repetitive firing was interrupted by silent periods (Figure 5B), or prolonged inter-spike intervals (ISIs), during which the FS cell entered a subthreshold state in the continuous presence of excitatory current (Golomb et al., 2007; Ermentrout and Wechselberger, 2009). This irregularity of FS cell firing was previously termed stuttering (Markram et al., 2004; Descalzo et al., 2005; Goldberg et al., 2008; Povysheva et al., 2008; Helm et al., 2013). To quantify the irregularity of firing, we calculated the coefficient of variation of the ISIs (CV_ISI_) in the spike trains (Povysheva et al., 2008; Helm et al., 2013). In the FS cell model with intact g_Kv2_, the CV_ISI_ diminished sharply as the applied current increased just above threshold (Figure 5C). In contrast, in the FS model with reduced g_Kv2_, larger applied currents were necessary to decrease stuttering and the CV_ISI_ (Figure 5C). Moreover, the CV_ISI_ decreased with increasing g_Kv2_ levels (Figure 5C,D), supporting the idea that Kv2.1 currents prevent stuttering.

**Figure 5.**
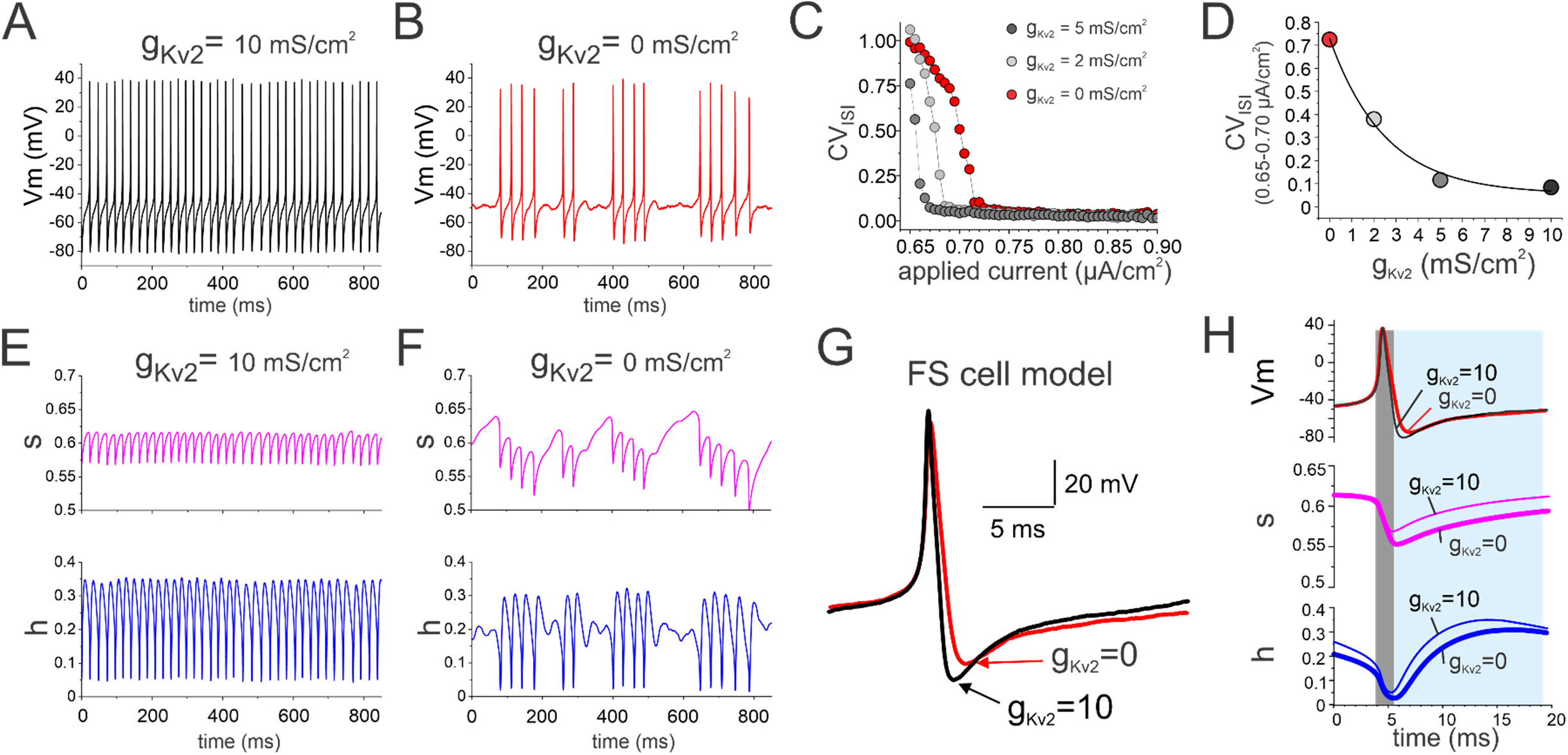
Simulations in a fast spiking (FS) cell model assessing the effect of reduced Kv2.1 conductance on the FS cell properties. **A)** Spike train evoked in the model FS cell with Kv2.1 conductance (g_Kv2_) intact. **B)** Spike train evoked in the model FS cell without g_Kv2_. **C)** Plots of the coefficient of variation of the inter spike interval (CV_ISI_) in spike trains as a function of applied current for the model FS cell with intact and reduced levels of g_Kv2_. **D)** Graph of CV_ISI_ in spike trains evoked by applied currents (averaged for 0.65-0.70 μA/cm^2^) in the model cell with intact and reduced levels of g_Kv2_. **E)** Time course of the Hodgkin-Huxley inactivation gating variables s (top) and h (bottom) for the FS cell model with intact Kv2, during the spike train shown in A). **F)** Time course of the Hodgkin-Huxley inactivation gating variables s (top) and h (bottom) for the FS cell model with reduced Kv2, during the spike train shown in B). **G)** Superimposed action potentials from the FS cell model with intact (black) and reduced (red) Kv2.1 The arrows indicate the maximal hyperpolarization during the afterhyperpolarization, which was smaller in the FS cell model with reduced Kv2.1. **H)** Superimposed action potentials (top), and simultaneous time courses of the inactivation gating variables s (middle) and h (bottom). The thin lines and thick represent, respectively, the data from the model with intact or reduced Kv2.1. The gray rectangle indicates the portion of the action potential where most of the inactivation of the Na^+^ conductance occurs. The light blue rectangle indicates the portion of the repolarization phase and afterhyperpolarization where most of the recovery from inactivation of the Na^+^ conductance takes place.

Next, we used the FS cell model to examine whether the strong stuttering phenotype produced by reduced levels of Kv2.1 conductance is associated with increased Na^+^ channel inactivation, since Kv2.1 current was reported to facilitate repetitive firing by reducing Na^+^ channel inactivation in other types of neurons (Guan et al., 2013; Romer et al., 2019). We assessed Na^+^ channel inactivation by plotting the time course of the Hodgkin-Huxley inactivation gating variables, s and h, throughout the spike trains. During continuous firing in the FS model with intact g_Kv2_, the Na^+^ conductance consistently inactivated during each spike, as indicated by downward shifts of s and h values, and recovered rapidly from inactivation during each ISI (Figure 5E). In the FS cell model with reduced g_Kv2_, the Na^+^ conductance inactivation was stronger, as revealed by a greater reduction in s and h values with each spike (Figure 5F). Moreover, during successive spikes the Na^+^ channel inactivation summated until the FS cell model with reduced g_Kv2_ became refractory, and resumed firing when the Na^+^ conductance recovered, producing the long ISIs characteristic of stuttering (Figure 5B and F). Our simulations therefore suggest that Kv2.1 currents facilitate repetitive firing in FS cells by preventing Na^+^ channel inactivation, and hence that loss of g_Kv2_ may produce stuttering secondary to enhanced and prolonged inactivation of Na^+^ channels.

Notably, in the FS cell model with reduced g_Kv2_, the afterhyperpolarization associated with single spikes had smaller amplitude (Figures 5G, see also Figure 5A,B), whereas the action potential repolarization was slower (Figure 5G). The inactivation of Na^+^ channels and their recovery from inactivation both are time- and voltage-dependent, while the recovery requires a hyperpolarized potential (Johnston and Wu, 1995), a feature that is well described in Hodgkin-Huxley models. Hence, the reduction of the afterhyperpolarization amplitude may contribute to the mechanisms by which decreasing Kv2.1 conductance increases Na^+^ channel inactivation, thus promoting stuttering. To investigate whether changes in the afterhyperpolarization properties contribute to greater Na^+^ channel inactivation, we examined the time course of inactivation around the time window of an action potential. The time window of Na^+^ channel inactivation by the action potential (gray rectangle in Figure 5H) starts during the ascending/depolarization phase, reaching maximal inactivation (lowest value of s or h) during the descending/repolarization phase, until recovery from inactivation begins. The time window when Na^+^ channels recover from inactivation (light blue rectangle in Figure 5H) started during the action potential descending/repolarization phase but largely coincided with the afterhyperpolarization. Hence, as a likely consequence of the slower action potential repolarization and smaller afterhyperpolarization amplitude, in the model with reduced g_Kv2_ the maximal inactivation of the Na^+^ conductance was larger and the recovery from inactivation was slower (Figure 5H), possibly explaining the temporal summation of inactivation during successive spikes (Figure 5F). Our simulations therefore suggest that greater inactivation and impaired recovery of Na^+^ channels from inactivation, secondary to reduced Kv2.1-mediated currents, affect repetitive firing in FS cells. This idea is consistent with experimental data showing that rapid recovery from inactivation of the Na^+^ conductance is essential for the FS phenotype of PV neurons (Martina and Jonas, 1997).

### *Kcns3* deficiency alters repetitive firing in PVBCs

The results of our simulations in the FS cell model suggest that if *Kcns3* deficiency decreases the Kv9.3-mediated amplification of the Kv2.1 current in PV neurons, it should increase stuttering. To test the predictions from the simulations, we assessed the firing properties of PVBCs in acute slices from the PFC of G42-*Kcns3*^+/+^ and G42-*Kcns3*^neo/neo^ mice. As observed in the FS model with low g_Kv2_, in PVBCs from G42-*Kcsn3*^neo/neo^ mice the action potential repolarization appeared slower and the afterhyperpolarization amplitude significantly smaller relative to G42-*Kcns3*^+/+^ mice (Figure 6A). Hence, *Kcns3* deficiency had similar effects to those produced by reducing Kv2.1 in the FS cell model.

**Figure 6.**
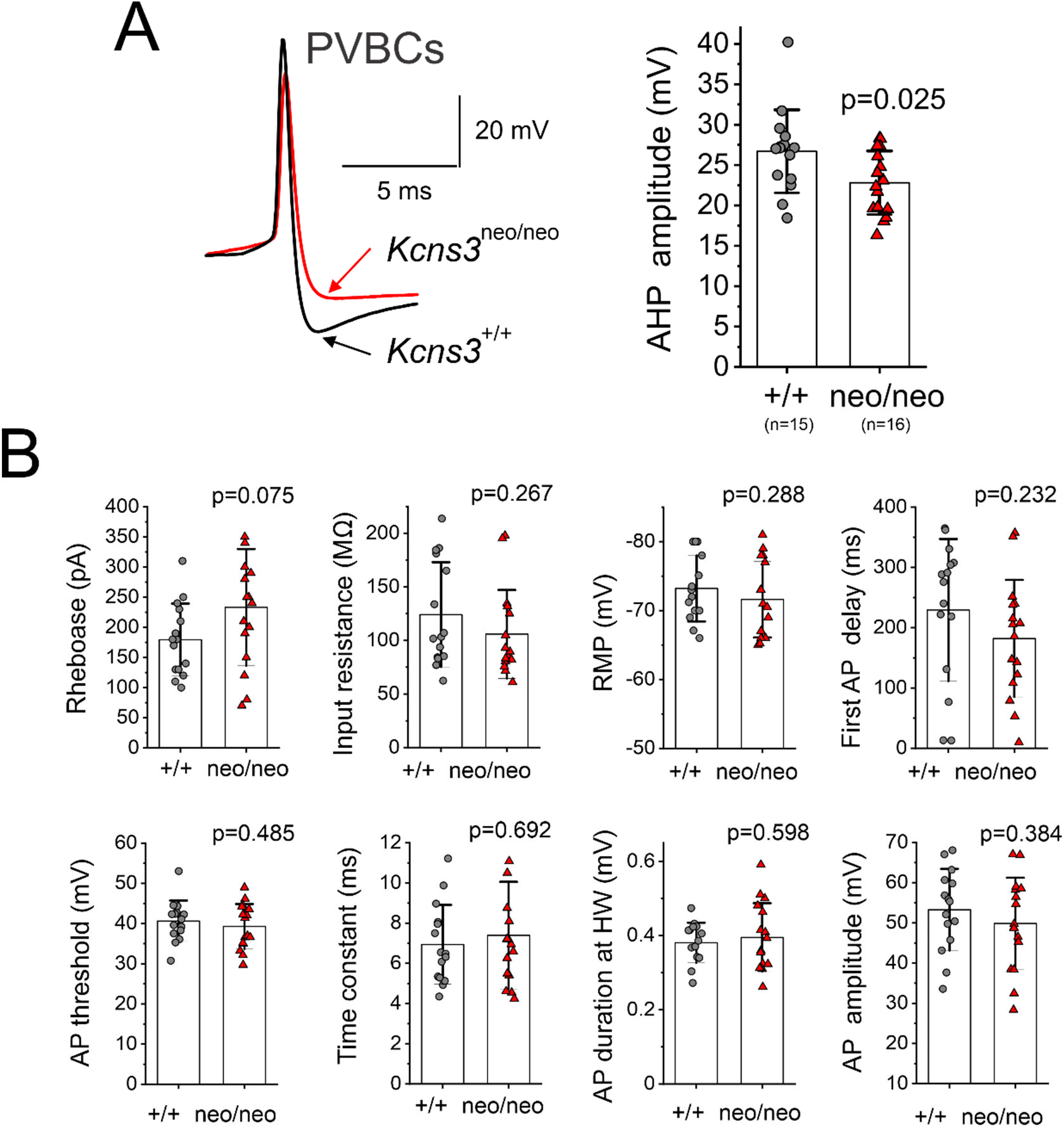
Effects of *Kcns3* deficiency on the membrane properties of parvalbumin-positive basket cells (PVBCs). **A)** Left panel, representative action potentials recorded from PVBCs from G42-*Kcns3*^+/+^ and G42-*Kcns3*^neo/neo^ mice. The arrows indicate the maximal hyperpolarization during the afterhyperpolarization, which was smaller in PVBCs from *Kcns3*-deficient mice. Right panel, bar graph summarizing the afterhyperpolarization values for PVBCs from control and *Kcns3*-deficient mice. Here and in B), shown are the p values from Student’s t-tests using the original data without Ln transformation, except when indicated below. **B)** Membrane properties of PVBCs. The measured parameters are defined as follows, Rheobase: current threshold to fire at least one action potential (AP); RMP: resting membrane potential; First AP delay: delay to fire the first AP with rheobase current stimulation; AP threshold: voltage threshold to fire an AP; Time constant: membrane time constant; AP amplitude: voltage between AP threshold and AP peak; AP duration at HW: width of the AP at half maximal amplitude. For RMP, the Shapiro-Wilk test p values were 0.031 for original data and 0.04 for Ln-transformed data, thus comparison was done using Mann-Whitney’s test. For Rin, the Shapiro-Wilk test p values were 0.002 for original data and 0.055 for Ln-transformed data. For the membrane time constant, Shapiro-Wilk test p values were 0.03 for original data and 0.63 for Ln-transformed data, thus differences between group means were tested using Student’s t test after Ln transformation.

We assessed the general membrane properties of PVBCs, which are essential for repetitive firing (Miyamae et al., 2017), in G42-*Kcns3*^+/+^ (15 PVBCs from 7 mice) and G42-*Kcns3*^neo/neo^ (16 PVBCs from 8 mice) mice. Except for the afterhyperpolarization (Figure 6A), the membrane properties did not differ by *Kcns3* genotype (Figure 6B). The threshold current necessary to evoke spikes, or rheobase, was ~50 pA higher in PVBCs from *Kcsn3*-deficient mice, although the difference was non-significant (Figure 6B). Moreover, the rheobase was highly variable (Miyamae et al., 2017), and individual PVBCs from either genotype displayed high or low rheobase (Figure 6B).

To test whether *Kcns3* deficiency affects repetitive firing, we elicited spike trains applying steps of depolarizing current in 10 pA increments (Figure 7). Most PVBCs from G42-*Kcns3*^+/+^ mice displayed the continuous firing with nearly constant ISIs (Figure 7A) classical of FS neurons (Goldberg et al., 2008). In contrast, many PVBCs from G42-*Kcns3*^neo/neo^ mice showed spike trains with strong stuttering (Figure 7B) which was very similar to that observed in the FS cell model with reduced g_Kv2_ (Figure 5B), and required relatively large stimulus currents to convert into continuous firing (Figure 7B).

**Figure 7.**
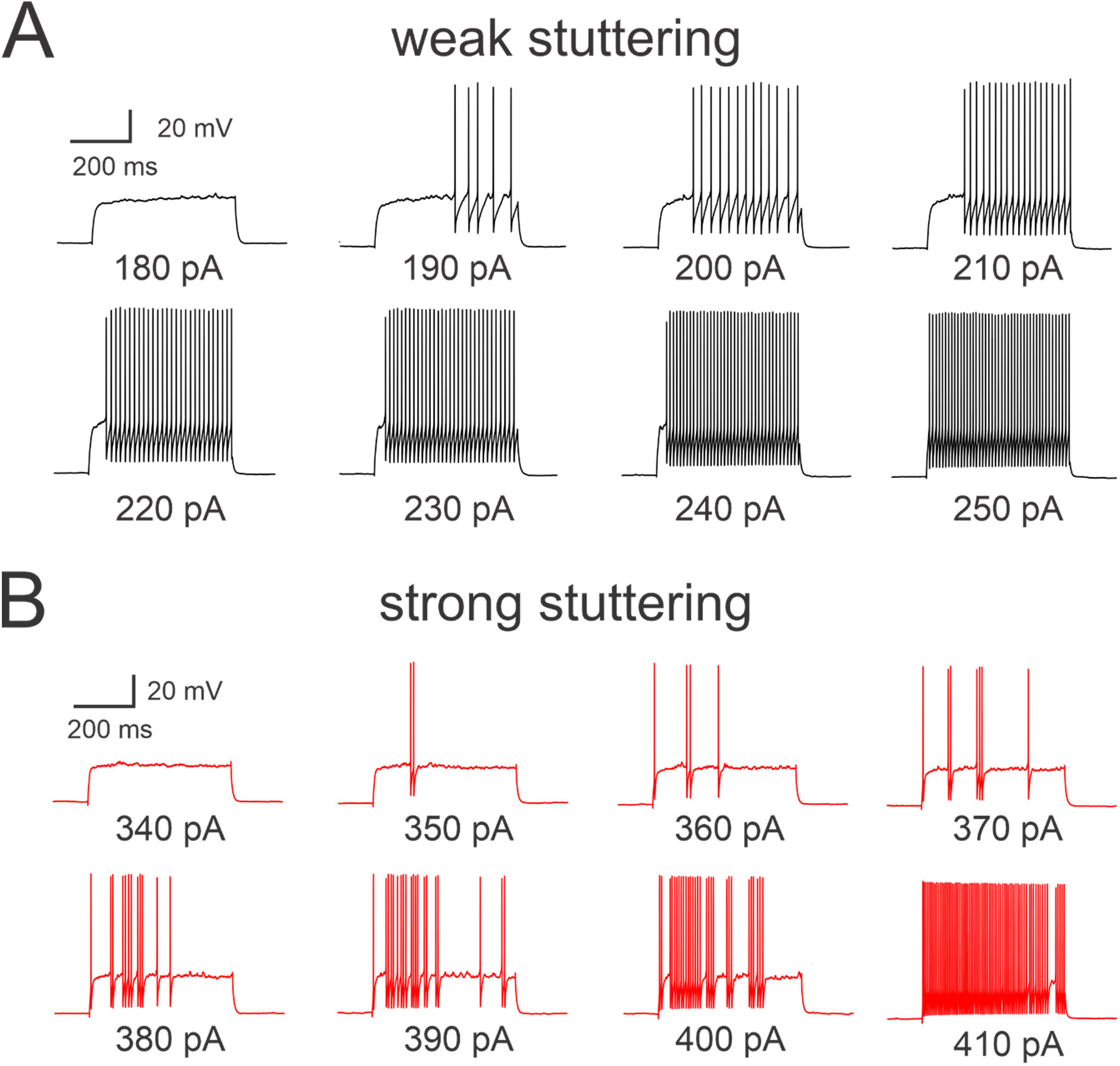
**A)** Representative spike trains evoked by injection of depolarizing current steps into a weak stuttering PVBC in the prefrontal cortex (PFC) of a G42-*Kcns3*^+/+^ mouse. **B)** Representative spike trains evoked by injection of depolarizing current steps into a PV cell with strong stuttering phenotype in PFC of a G42-*Kcns3*^neo/neo^ mouse.

Raster plots and plots of the CV_ISI_ built from the spike trains showed marked differences between PVBCs with weak or strong stuttering (Figure 8A,B). In weak stuttering PVBCs, the CV_ISI_ decreased sharply with stimulus currents near threshold (Figure 8A). In contrast, in strong stuttering PVBCs the CV_ISI_ was high near threshold and decreased progressively with stimuli well above threshold (Figure 8B). Most PVBCs from G42-*Kcns3*^+/+^ mice had weak stuttering (Figure 8C, left). In contrast, most PVBCs from G42-*Kcns3*^neo/neo^ mice showed strong stuttering (Figure 8C, right), and displayed significantly higher CV_ISI_ across a range of stimulus intensities (Figure 8D), as in the FS cell model with low g_Kv2_ (Figure 5C,D). These alterations of repetitive firing in PVBCs from *Kcns3*-deficient mice are similar to those predicted by our simulations in the FS cell model.

**Figure 8.**
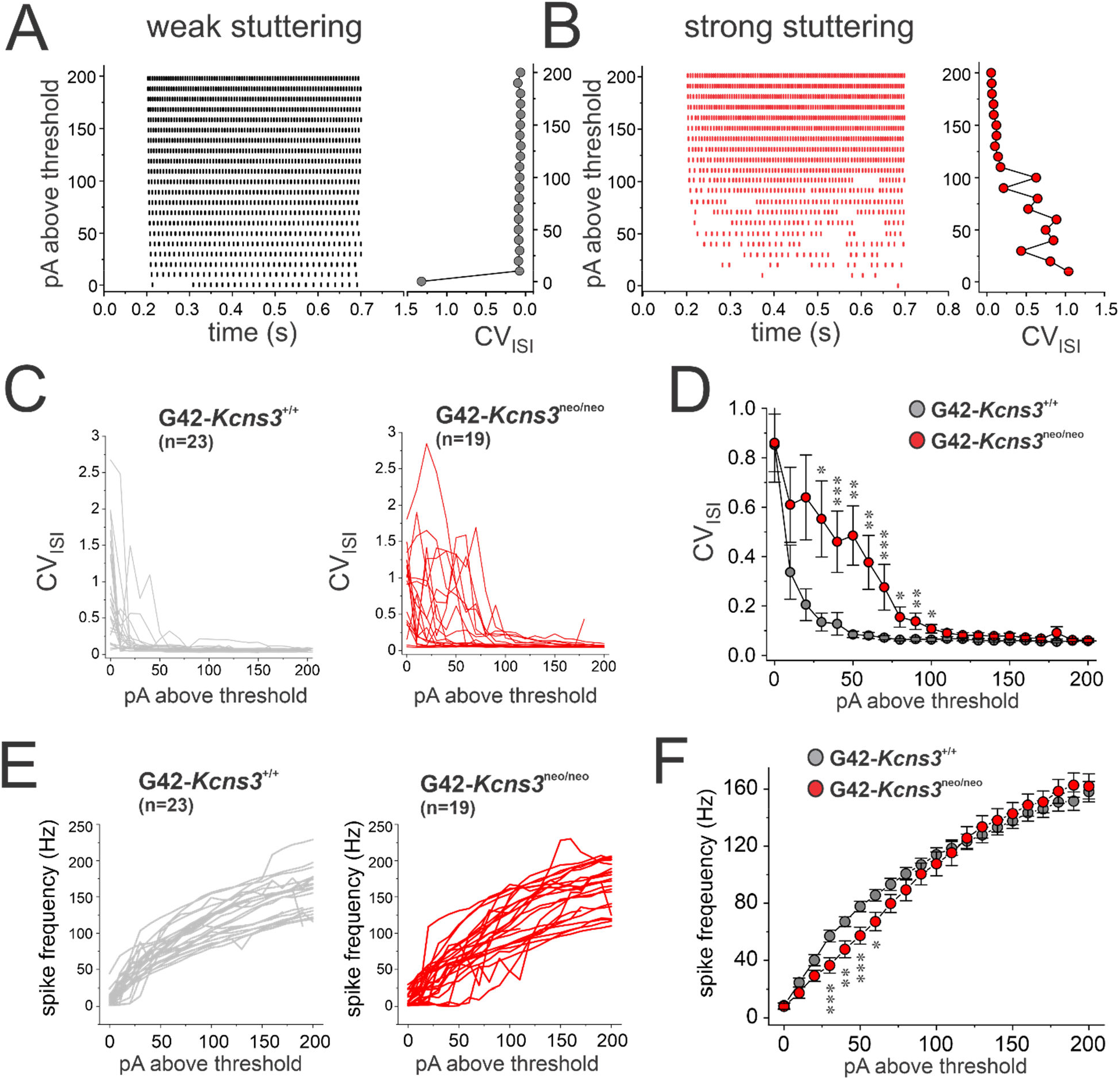
Effects of *Kcns3* deficiency on the variability of inter spike intervals (ISIs) in spike trains evoked by injection of current steps. **A)** Left panel, representative raster plot for spike trains evoked in a weak stuttering parvalbumin-positive basket cell (PVBC) in the prefrontal cortex (PFC) of a G42-*Kcns3*^+/+^ mouse. Right panel, coefficient of variation of ISIs (CV_ISI_) (X-axis) graphed as a function of stimulus current above threshold (Y-axis). **B)** Representative raster plot and CV_ISI_ graph as in A), but for spike trains evoked in a strong stuttering PVBC in PFC of a G42-*Kcns3*^neo/neo^ mouse. **C)** Graphs of CV_ISI_ as a function of stimulus current above threshold for each of the PVBCs of G42-*Kcns3*^+/+^ (left, n=23 cells from 11 mice) and G42-*Kcns3*^neo/neo^ mice (right, n=19 cells from 8 mice). **D)** Graphs of CV_ISI_ (mean±SEM) as a function of stimulus current above threshold for the PVBCs of G42-*Kcns3*^+/+^ and G42-*Kcns3*^neo/neo^ mice displayed in C). Shapiro-Wilk tests of the original data produced p values <0.05 at all current levels 0-130pA above threshold and p values <0.05 with Ln-transformed data at all current levels above threshold except 10, 20, 30 and 40 pA. Hence, we employed Mann-Whitney’s tests. *: p<0.02, **: p<0.01, ***: p<0.005. **E)** Left, graphs of mean spike frequency per current step as a function of stimulus current above threshold for each of the PVBCs of G42-*Kcns3*^+/+^(left, n=23 cells from 11 mice) and G42-*Kcns3*^neo/neo^ (right, n=19 cells from 8 mice) mice. **F)** Graphs of spike frequency (mean±SEM) as a function of stimulus current above threshold for PVBCs of G42-*kcns3*^+/+^ and G42-*kcns3*^neo/neo^ mice. For most comparisons at current levels 0-100 pA above threshold, Shapiro-Wilk tests produced p values <0.05 with original and Ln-transformed data, hence Mann-Whitney’s tests were employed. *: p<0.02, **: p<0.01, ***: p<0.005.

In strong stuttering PVBCs the spike trains had a larger proportion of long ISIs (Figure 7B), suggesting that stuttering depresses PVBC spike output. To assess whether stuttering significantly affects PVBC spike output, we built frequency-current (f-I) plots. The individual (Figure 8E) and average (Figure 8F) f-I plots showed lower spike output in G42-*Kcns3*^neo/neo^ mice with currents ≥ 20 pA and ≤ 70 pA above threshold, a stimulus range that elicited spiking in the gamma frequency band (30-80 Hz).

In other types of neurons, Kv2.1 currents control spike frequency adaptation (Guan et al., 2013; Romer et al., 2019), suggesting that Kv2.1-Kv9.3 channels may contribute to the low spike frequency adaptation typical of FS cells (Hu et al., 2014). Therefore, in spike trains elicited by current steps in the experiments depicted in Figure 8, we assessed spike frequency adaptation by computing the ISI_ratio_, or ratio between the last and first ISI, which is expected to produce values near 1 with low spike frequency adaptation. As shown in Table 2, at the stimulus range in which *Kcns3* deficiency increased stuttering (<100 pA above threshold) the ISI_ratio_ did not differ between genotypes. In contrast, with strong stimuli (>100 pA above threshold), that elicited firing at frequency>120 Hz while PVBCs from G42-*Kcns3*^+/+^ mice showed an ISI_ratio_ ~1, PVBCs from G42-*Kcns*3^neo/neo^ mice displayed significantly higher (~25-30%) ISI_ratio_ (Table 2). These data suggest that Kv2.1-Kv9.3 channels regulate spike frequency adaptation exclusively in a strong stimulus range where the CV_ISI_ was not significantly affected. Conversely, the ISI_ratio_ was not affected by *Kcns3* deficiency within the low stimulus range where the CV_ISI_ was significantly altered. Hence, the effects of *Kcns3* deficiency on the CV_ISI_ and the ISI_ratio_ may be mediated by different mechanisms. Moreover, the effects on the CV_ISI_ likely have a greater impact on recruitment of PVBCs by natural stimuli, since the changes in ISI_ratio_ were observed only with very strong stimuli.

**Table 2.** Effects of *Kcns3* deficiency on the adaptation ratio measured during spike trains elicited by current steps in PVBCs.

|  | G42-*Kcns3*<sup>+/+</sup><br>(23 cells, 11 mice) | G42- *Kcns3*<sup>neo/neo</sup><br>(19 cells, 8 mice) | p value |
| --- | --- | --- | --- |
| ISI<sub>ratio</sub> at 50 pA | 1.03±0.18 | 1.12±0.71 | p=0.591 |
| ISI<sub>ratio</sub> at 80 pA | 1.06±0.23 | 1.16±0.27 | p=0.208 |
| ISI<sub>ratio</sub> at 100 pA | 1.01±0.24 | 1.25±0.26 | p=0.003 |
| ISI<sub>ratio</sub> at 150 pA | 1.04±0.26 | 1.31±0.20 | p=0.002 |

Shapiro-Wilk and D-Agostino K square normality tests rejected a normal distribution of the residuals or the residuals of the Ln-transformed data for the ISI ratio with 80, 100 and 150 pA stimuli. Hence comparisons were performed using Mann-Whitney tests.

### *Kcns3* deficiency disrupts PV neuron spike output during UP states

Our experiments and simulations using standard steps of depolarizing current showed that *Kcns3* deficiency alters the classical FS phenotype of PV neurons, producing irregular firing, or stuttering, and decreasing spike output in the gamma frequency band. Current steps, however, may not reveal the effects of *Kcns3* deficiency on PV neuron activity driven by more natural stimuli. Thus, next we assessed the effects of *Kcns3* deficiency on the transformation of excitatory input into spike output, studying spike trains elicited by injecting excitatory current with the shape of synaptic input observed during cortical UP states (Figure 9A). UP states are episodes of increased recurrent network activity that involve pyramidal cells and PV neurons (Puig et al., 2008; Fanselow and Connors, 2010; Zucca et al., 2017), which depolarize single neurons near spike threshold (Shu et al., 2003; Hasenstaub et al., 2005). To investigate the effects of *Kcns3* deficiency on PVBC spike output during UP states, we first induced UP states in PFC slices using a previously described protocol (Sanchez-Vives and McCormick, 2000). Synaptic input observed in single neurons during UP states was recorded, stored digitally and used for stimulation with current injection, comparing the response of PVBCs from *Kcns3^+/+^*versus *Kcns3*^neo/neo^ mice. We injected the UP state-like currents increasing, in 10 pA increments, the mean current injected during the first UP state (UP_1_) until UP_1_ elicited spikes, defining stimulus strength 1, and up to 4 stimulus strengths.

**Figure 9.**
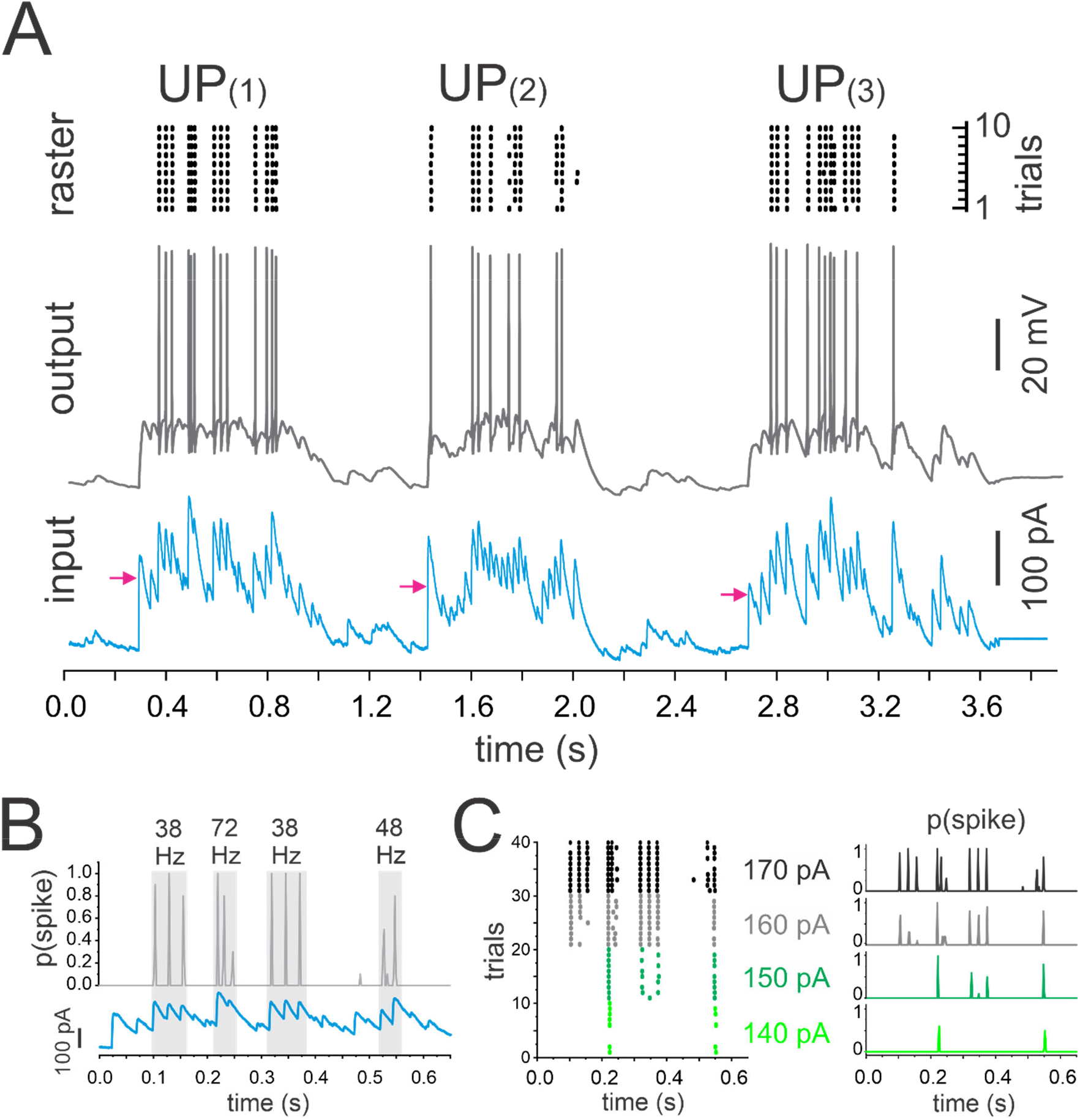
The current injection paradigm mimicking UP states, employed to assess the effects of *Kcns3* deficiency. **A)** The traces at the bottom (input) show the injected depolarizing current mimicking synaptic input during three UP states (UP_(1)_, UP_(2)_, and UP_(3)_) with two intermediate DOWN states. The arrows represent the mean injected current obtained by averaging the current injected across the duration of each UP state. Traces in the middle (output) show the spike output evoked in a parvalbumin-positive basket cell (PVBC) by the UP state current injection. The raster plots at the top were obtained from the spike trains evoked in the PVBC by 10 repetitions of the UP state current injection at stimulus strength 4 in 10 consecutive trails. **B)** The top graph shows the average spike probability, p(spike), computed in 2 ms bins throughout the injection of UP_(1)_ repeated ten times at the stimulus strength 4. The bottom graph shows the depolarizing current injected during UP_(1)_. The gray rectangles indicate the time windows of bursts of gamma band activity, with frequency measured from the intervals between subsequent peaks of p(spike). **C)** Left, raster plots from spike trains evoked by injection into a PVBC of UP_(1)_ at stimulus strengths 1 to 4 (the mean injected current during UP_(1)_ was 140, 150, 160 and 170 pA in this example). Right, spike probability p(spike) computed from the raster plots in the left panel.

During UP states, PVBC spikes were elicited at the peaks of injected current (Figure 9B) as expected with injections of variable current (Mainen and Sejnowski, 1995). We assessed the response of PVBCs to the UP state-like stimulus computing the spike probability, p(spike), in 2 ms windows throughout the spike trains elicited by ten identical repeats of each stimulus pattern and strength (Figure 9C). The curves of p(spike) showed peaks time-locked to the peaks of injected current (Figure 9B), while the height of the peaks of p(spike) increased with stimulus strength (Figure 9C), reflecting the increased likelihood of recruiting PVBCs as the input strength increased.

Consistent with previous reports that UP states are associated with gamma frequency (30-80 Hz) network activity (Shu et al., 2003; Hasenstaub et al., 2005), the synaptic activity recorded during UP states showed successive peaks at gamma frequency (Figure 9A,B). When injected into PVBCs, the UP state-like stimuli produced brief bursts of spikes at gamma frequency (Figure 9B). Therefore, the injection of UP state-like currents allowed us to test whether *Kcns3* deficiency affects the response to natural inputs that elicit PVBC activity in the gamma frequency band.

To compare between genotypes the spike output during UP states, we obtained a curve of average p(spike) by averaging across neurons the curves obtained for each neuron at a given stimulus strength (Figure 9B), in the PVBCs from G42-*Kcns3*^+/+^ (n=25 cells from 15 mice), and from G42-*Kcns3*^neo/neo^ mice (n=25 cells from 14 mice). The curves of average p(spike) showed peaks with identical timing in PVBCs from G42-*Kcns3*^+/+^ and G42-*Kcns3*^neo/neo^ mice, as the PVBCs from either genotype responded to identical peaks of input current (Figure 10A). The peaks of average p(spike) with identical timing share, across genotypes, the previous history of variable input current and hence were paired for comparison between genotypes. We found that in most pairs of average p(spike) peaks, the peak in the PVBCs from *Kcns3*-deficient mice had lower height (Figure 10A,B). Pair-wise comparisons (Figure 10B) of the differences in the height of the paired peaks showed that the reduced height of the peaks of average p(spike) became significant as the stimulus strength increased (Figure 10C). These data suggest that *Kcns3* deficiency reduces the probability of PVBC recruitment by UP state-like stimulus patterns.

**Figure 10.**
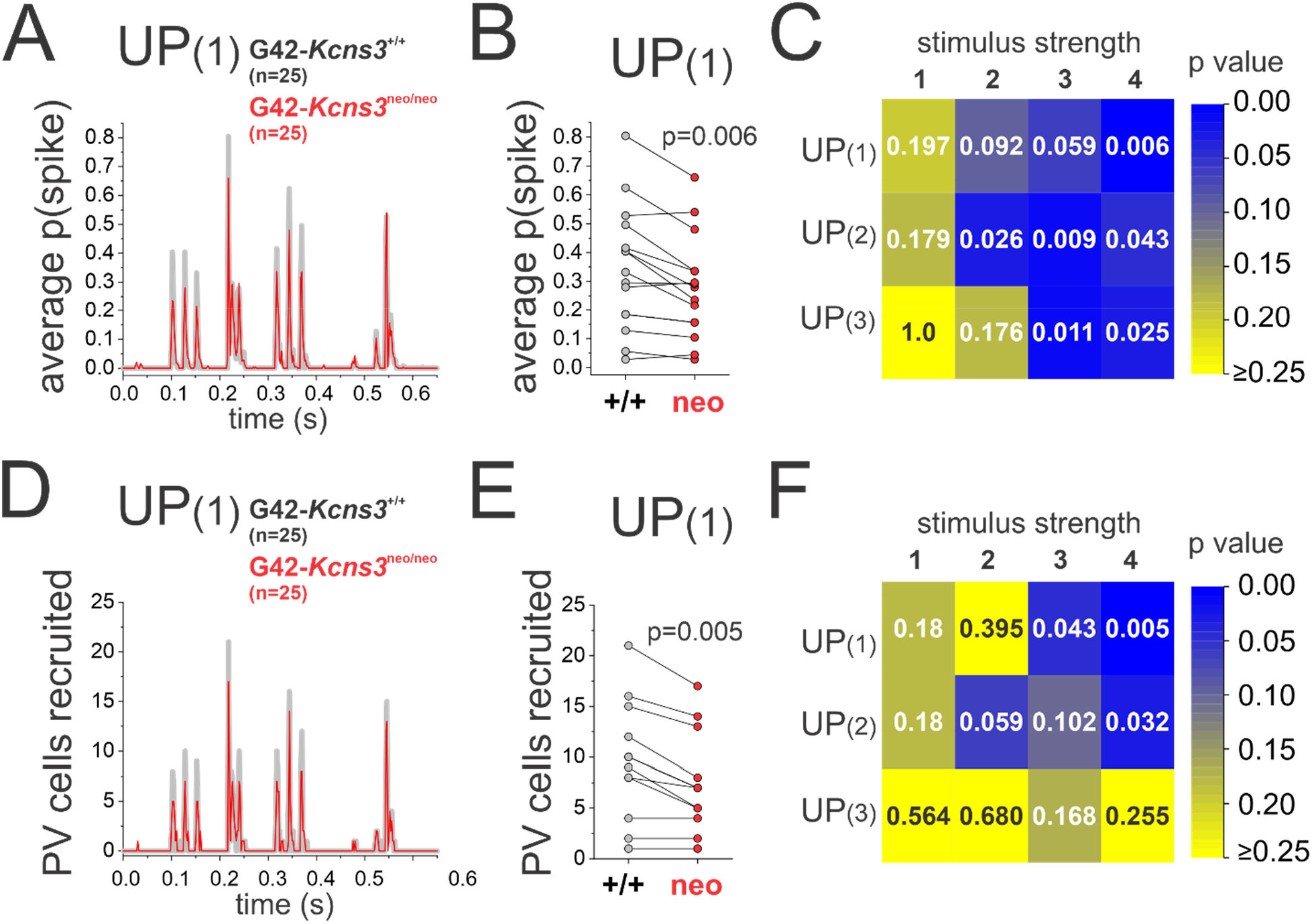
Effects of *Kcns3* deficiency on the spike output of parvalbumin-positive basket cells (PVBCs) during UP states. **A)** Curves of mean spike probability, p(spike), during UP_(1)_ at stimulus strength 4, computed by averaging p(spike) across 25 PVBCs neurons from 15 G42-*Kcns3*^+/+^ mice and 25 PV neurons from 14 G42-*Kcns3*^neo/neo^ mice. **B)** Paired comparison of p(spike) for the corresponding peaks of spike probability depicted in C. **C)** Matrix plot showing the p values of paired comparisons for matching peaks of p(spike) between *G42-kcns3^+/+^* and G42-*kcns3*^neo/neo^ mice during injection of UP_(1)_, UP_(2)_ and UP_(3)_ at stimulus strengths 1-4. For most combinations of UP state and stimulus strength, Shapiro-Wilk analysis produced p values <0.05 with the original and the Ln-transformed data. Hence, here and in B, we report the p values of Wilcoxon Signed Rank tests. **D)** Curves of the number of PVBCs recruited by a single presentation of UP_(1)_ at stimulus strength 4, computed for the 25 PV neurons from G42-*Kcns3*^+/+^ and 25 cells from G42-*Kcns3*^neo/neo^ mice. **E)** Paired comparison of the number of PVBCs recruited by a single presentation of UP_(1)_ for the matching peaks depicted in D. **F)** Matrix plot showing the p values of paired comparisons for matching peaks of number of PVBCs recruited among the 25 PV neurons between G42-*Kcns3*^+/+^ and G42-*Kcns3*^neo/neo^ mice during injection of UP_(1)_, UP_(2)_ and UP_(3)_ at stimulus strengths 1-4. For most combinations of UP state and stimulus strength, Shapiro-Wilk analysis produced p values <0.05 with the original and the Ln-transformed data. Hence, here and in E, we report the p values of Wilcoxon Signed Rank tests.

During natural network activity, the patterns of input associated with discrete UP states are likely to be unique to each UP state. Hence, we wondered whether the changes in average p(spike) revealed using 10 identical repeats of each stimulus pattern may predict the changes in the probability of recruiting PVBCs with single applications of a stimulus pattern. If so, then *Kcns3* deficiency should reduce the number of PVBCs recruited by the first of the ten repeats of each UP state and stimulus strength. The number of PVBCs recruited by the first repeat of each UP state-like stimulus showed peaks that matched in timing the peaks of average p(spike) (Figure 10D). The peaks in the number of recruited PVBCs were lower in *Kcns3*-deficient mice (Figure 10E), and the reduction in the numbers of PVBCs recruited became significant with increasing stimulus strength (Figure 10F). These data thus suggest that *Kcns3* deficiency reduces the fraction of PVBCs recruited when the local excitatory network generates UP state activity particularly, but not exclusively, in the gamma frequency band.

## Discussion

We assessed the role of Kv9.3 subunits in PV neuron electrophysiology using a line of *Kcns3*-deficient mice. Contrary to our prediction that *Kcns3* deficiency would alter the EPSP decay time course (Georgiev et al., 2012; Georgiev et al., 2014), we found that *Kcns3* deficiency did not alter sEPSPs, suggesting that the small depolarization elicited by EPSPs is not sufficient to activate Kv2.1-Kv9.3 channels. Given that Kv9.3 subunits amplify the Kv2.1 current, and that Kv2.1 currents regulate repetitive firing in other types of neurons, we hypothesized that lower Kv9.3 subunit levels may affect repetitive firing in PV neurons. We tested the effects of decreasing Kv2.1 on repetitive firing in a computational model of PV FS cells. Our simulations showed that reducing Kv2.1 produces irregular ISIs, or stuttering, via an increase of Na^+^ channel inactivation possibly caused by a smaller afterhyperpolarization amplitude. Consistent with our simulations, PVBCs from *Kcns3*-deficient mice had smaller afterhyperpolarization amplitude and exhibited strong stuttering, indicating that *Kcns3* deficiency critically affects the classical FS phenotype of PVBCs. Disruption of the FS phenotype was accompanied by impaired recruitment of PVBCs by natural patterns of input such as those observed during cortical UP states, which normally recruit PV neuron firing at gamma frequency band. Our experiments with *Kcsn3*-deficient mice suggest specific ways in which *KCNS3* deficiency in schizophrenia might affect PVBC and cortical circuit function.

### Gene expression changes in *Kcns3*^neo/neo^ mice

The insertion of the trapping cassette into the intron directly upstream of the protein coding exon of the *Kcns3* gene (Figure 1A), resulted in a partial 45% reduction of *Kcns3* mRNA in cortical tissue from Kcns3^neo/neo^ mice, as revealed by qPCR using primers that amplified a protein-coding sequence of the mRNA (Figure 1). Such partial effects of trapping cassettes have been previously reported for multiple genes (Serafini et al., 1996; Yeo et al., 1997; Voss et al., 1998) and attributed to the splicing of pre-mRNAs around a trapping cassette or to transcription driven by an alternative promoter. Consistent with the qPCR results, single-cell ISH analysis showed a 50% decrease of *Kcns3* mRNA expression in cortical PV neurons of *Kcns3*^neo/neo^ mice (Figure 2). The distribution of *kcns3* mRNA expression per single PV cell partially overlapped between *Kcns3^+/+^* and *Kcns3*^neo/neo^ mice, showing that some PV neurons in *Kcns3*^neo/neo^ mice had *Kcns3* mRNA levels typical of *Kcns3^+/+^* mice and thus may display an intact physiological phenotype. Consistent with this interpretation, some PV neurons from *Kcns3*^neo/neo^ mice had unaltered electrophysiology (Figure 8).

Kv9.3 subunits form heteromeric Kv2.1-Kv9.3 channels. Because expression of *Kcnb1*,the gene encoding Kv2.1, was not altered in PV neurons from *Kcns3^neo/neo^* mice (Figure 2), the observed alterations of PVBC physiology in *Kcns3^neo/neo^* mice are likely specific to lower levels of Kv9.3 subunits.

### *Kcns3* deficiency does not affect synaptic input onto PV neurons

We proposed previously that Kv2.1-Kv9.3 channels are more effectively activated by subthreshold EPSPs than homomeric Kv2.1 channels (Georgiev et al., 2012; Georgiev et al., 2014), and accelerate the EPSP decay. However, we found that sEPSPs, including the sEPSPs >1mV, were unaltered in PVBCs from *Kcns3*-deficient mice (Figure 4). One possibility is that due to their voltage dependence of activation (Bocksteins, 2016), Kv2.1-Kv9.3 channels are minimally activated by EPSPs. Due to severe attenuation during propagation to the soma EPSPs recorded at the PV neuron soma could have amplitude up to ~20 times smaller than the EPSPs generated in the dendrites (Hu et al., 2010; Norenberg et al., 2010). Interestingly, analysis of the subcellular distribution of Kv2.1 subunits in PV neurons suggests that Kv2.1-Kv9.3 channels are present in dendrites (Du et al., 1998). Hence, it is unlikely that the absence of Kv2.1-Kv9.3 channel-mediated regulation of the sEPSP shape is due to the small amplitude of subthreshold EPSPs recorded at the soma. Our data therefore suggest that depolarizations larger than somatic or dendritic EPSPs, such as those produced by action potentials, may be necessary to activate Kv2.1-Kv9.3 channels.

### PV neurons from *Kcns3*-deficient mice have disrupted input-output transformation properties

To assess how *Kcns3* deficiency could affect the FS phenotype of PV neurons, we used a computational FS cell model to test the effects of decreasing the Kv2.1 conductance, the effect expected after the loss of Kv9.3-mediated amplification of the Kv2.1 current. We found that in the FS cell model with reduced Kv2.1 conductance, spike trains displayed strong stuttering (Figure 5). Our simulations also suggest that one of the mechanisms by which reduced Kv2.1 conductance may increase stuttering involves greater inactivation of the Na^+^ channels, possibly caused by a smaller afterhyperpolarization amplitude (Figure 5). Our simulation data therefore lead us to predict that due to lower Kv9.3 subunit levels, PVBCs from G42-*Kcns3*^neo/neo^ mice should display strong stuttering. In experiments with G42-*Kcns3*^neo/neo^ mice, PVBCs had both smaller afterhyperpolarization and strong stuttering (Figures 6-8), in a manner consistent with the predictions of our simulations in the FS cell model.

The experiments injecting current steps also showed that the increased stuttering reduces the spike output in the gamma frequency band (Figure 8). Moreover, *Kcns3* deficiency also decreased PVBC spike output produced by stimulation with UP state-like stimulus currents (Figure 9). UP states are episodes of increased network activity that engage pyramidal cells and PV neurons (Puig et al., 2008; Fanselow and Connors, 2010; Zucca et al., 2017), and produce rhythmic network activity in the gamma band (Shu et al., 2003; Hasenstaub et al., 2005). We found that PVBCs recruited by the UP state-like stimulus fired brief bursts of spikes at gamma frequency (Figure 9), and that the probability of PVBC firing during these gamma bursts was reduced in G42-*Kcns3^neo/neo^* mice (Figure 10). Our data therefore suggest that Kv9.3 channels critically shape PVBC electrophysiology tested using standard (current steps) or more natural (UP state-like) stimuli. Moreover, our data indicate that *Kcns3* deficiency disrupts the capacity of a network of PVBCs to fire during UP state episodes that normally produce bursts of gamma frequency firing.

### Implications for cortical circuit dysfunction in schizophrenia

The variety of Kv channel genes expressed by PV neurons is highly conserved between mouse and human neocortex (Okaty et al., 2009; Enwright et al., 2018; Krienen et al., 2019)(http://interneuron.mccarrolllab.org/). These and other similarities in voltage-gated channel gene expression suggest that PV neurons from human neocortex display FS properties, as verified experimentally (Wang et al., 2016; Stedehouder et al., 2017; Szegedi et al., 2017; Stedehouder et al., 2019; Szegedi et al., 2020). Hence, *Kcns3* deficiency may produce qualitatively similar electrophysiological changes in PV neurons whether from mouse or human cortex.

Previous postmortem studies reported that *KCNS3* mRNA levels were lower by 30-40% in PV neurons from the PFC of schizophrenia subjects compared with unaffected comparison subjects (Georgiev et al., 2014; Enwright et al., 2018). Therefore, our *Kcns3*-deficient mice with the 50% reduction in *Kcns3* mRNA levels in PV neurons appear to be an appropriate animal model to address the effects of the partial deficit of *KCNS3* expression in PV neurons observed in schizophrenia. Importantly, whereas *Kcnb1* mRNA levels were unaltered in *Kcns3*-deficient mice, *KCNB1* mRNA levels were lower in DLPFC PV neurons in schizophrenia in parallel with *KCNS3* mRNA levels (Georgiev et al., 2014), suggesting a greater reduction of Kv2.1 current in schizophrenia. In addition, comprehensive gene expression analysis (Enwright et al., 2018) demonstrated that Kv9.3 and Kv2.1 are the only Kv subunits with lower expression in PV neurons in schizophrenia. Hence, the physiological deficits in PV neurons might be qualitatively similar but quantitatively stronger in schizophrenia than in our line of *Kcns3*-deficient mice.

The mechanisms leading to lower *KCNS3* expression in schizophrenia are currently unknown. Remarkably, the expression of Kv channel genes, including *Kcns3*, is activity-dependent (Dehorter et al., 2015; Lee et al., 2015), suggesting that downregulation of *KCNS3* expression in schizophrenia is secondary to the lower levels of cortical network activity (Lewis et al., 2012; Arion et al., 2015). Our experiments in *Kcns3*-deficient mice suggest that *KCNS3*downregulation in schizophrenia, although possibly triggered by reduced network activity, is not a homeostatic adaptation to restore PV cell activity. Instead, *Kcns3* downregulation seems to produce a deficit in PV cell recruitment.

In schizophrenia, the alterations in PV neurons due to *KCNS3* deficiency might interact with a decrease in excitatory input (Chung et al., 2016), additionally impairing their recruitment. Whereas impaired PVBC recruitment due to lower number of excitatory synapses is independent of the firing rate of presynaptic excitatory neurons, our experiments with *Kcns3*-deficient mice suggest that PVBC recruitment may be affected particularly, but not exclusively, during rhythmic excitatory synaptic input such as that observed during gamma band oscillations (Mann et al., 2005; Oren et al., 2006; Atallah and Scanziani, 2009). Thus, *KCNS3* deficiency, together with other alterations in PV neurons (Dienel and Lewis, 2019), might contribute to the decrease in gamma oscillation power observed in schizophrenia under task conditions, such as those engaging working memory (Uhlhaas and Singer, 2015), that normally induce gamma band activity (Miller et al., 2018).

## Acknowledgments and Disclosures

This work was supported by the Japan Society for the Promotion of Science Grants-in-Aid 25293247 and 15H01280 to TH, and National Institutes of Health Grant Nos. MH51234 (to DAL) and P50MH103204 (to DAL, GG-B).

We thank Junko Konishi for the maintenance of mouse colonies, Olga Krimer for assistance with brain slice preparation, tissue staining and digital reconstructions of neuron morphology, and Yelena Gulchina for electrophysiological experiments during early stages of this project.

DAL currently receives investigator-initiated research support from Merck and Pfizer and serves as a consultant to Astellas. The other authors report no biomedical financial interests or potential conflicts of interest.

